# Size-dependent nucleus-vacuole interactions in budding yeast demonstrate a role for steric packing in organelle shape and positioning

**DOI:** 10.64898/2026.05.08.723889

**Authors:** Mary Mirvis, Oghosa H. Akenuwa, Irene Tan, Christopher T. Lee, Wallace F. Marshall

## Abstract

Although organelles are often studied one at a time, whole-cell imaging studies show that organelles take up a large part of the cell volume such that they are crowded together. Here we use whole cell soft X-ray tomography imaging to investigate how such crowding affects organelle size scaling, position, and shape, focusing on the nucleus and vacuole of budding yeast. We find that, as the vacuole enlarges beyond normal range, the nucleus loses its normal scaling relation with respect to cell volume, becomes displaced from its normal position near the cell center, and becomes progressively deformed from a sphere into a pancake shape. Using a whole-cell integrated shape modeling framework, we find that these changes are statistically correlated and give rise to distinct modes in cell organization space. Using a simplified mechanical model for two initially spherical compartments contained inside a confined intracellular space, we are able to recapitulate the effects seen in the experimental data, indicating that these observations are consistent with a purely mechanical interaction. Taken together, our work indicates that, in addition to the well-known protein-based organelle-organelle interactions, physical steric packing of organelles inside a limited cellular volume also plays a large role in the inter-organelle relationships and the overall geometry of the cell.

## Introduction

Physical density in the cellular context has come into focus as an important property of cells that can influence cellular mechanics, transport and diffusion dynamics, signaling and information transmission, stress response, and other crucial functions. At the biomolecular scale, molecular crowding in the cytoplasm and within subcellular compartments plays an important role in biochemical reaction rates and cellular homeostasis (Huang & Ferrell, 2026; Serrano et al., 2026). Mechanisms for maintaining and locally regulating molecular density have been uncovered. Cytoplasmic density is regulated to stay within a functional range, otherwise resulting in impacts on translation (Marini et al., 2020), cytoskeletal assembly dynamics (Garner et al., 2023; Molines et al., 2022), and formation of biomolecular condensates (van Tartwijk & Kaminski, 2022). At the tissue level, cellular crowding in tissues is known to be important for tissue mechanical properties, development, and cell signaling.

Between these scales, crowding in the mesoscale of membrane-bound organelles has been less studied. Classical electron micrographs visually demonstrate a high density of membranes, in addition to other mesoscale structures including cytoskeleton and other protein assemblies within the cytoplasm in small sections. Many observations of organelle interactions demonstrate that membranes of apposing organelles fit together hand-in-glove, matching their local curvatures (Friedman et al., 2011; Matos-Perdomo et al., 2022; Uchida et al., 2009), suggesting that the organelles may be packed closely enough together that they can affect each other’s shape.

Why might crowding or packing at the organelle scale be important? All cell types exhibit some degree of morphological heterogeneity, but there is typically a characteristic pattern to the positioning, shape, abundance, and size of organelles throughout the cell, which we refer to as cell anatomy. Anatomy may be significantly remodeled during cell state changes, such as during differentiation, cell division, metabolic shifts, and stress responses, but typically these remodeling events also occur in a stereotyped way. The robustness of anatomical patterns within a given cell type and context suggests the existence of active as well as passive, physical factors constraining organelle morphologies and higher order organization. While much of the focus on organelle shape determination has traditionally focused on membrane curvature-inducing proteins (Westrate et al., 2015), it seems likely that the inherent crowdedness of organelle membranes is an additional physical mechanism that could impact shape.

At the whole-organelle and whole-cell level, how crowded are intracellular membranes? How is this crowding regulated, and how does it factor into organelle structure, size control, biogenesis, function, and communication between organelles? To differentiate spatial constraints on this scale from that of molecular and tissue crowding, and to reflect the discrete nature of membrane-bound compartments we refer to this phenomenon as the “packing” of organelles or membranes. The overall degree and heterogeneity of crowdedness, and the effects of organelle packing, are difficult to ascertain without whole-cell 3D reconstructions of cells. Such imaging of whole cell anatomy is technically challenging to produce, requiring high 3D resolution imaging of a large volume, the ability to distinguish between multiple organelles, and segmentation methods that either rely on slow manual tracing or automated tools requiring extensive validation (Ekman et al., 2020; Erozan et al., 2024). A recent effort to curate whole-cell reconstruction data provides a view of anatomical patterns across eukaryotic cell types in studies that are generally small (Mirvis et al., 2026). The ability to visualize full volumetric maps of organelles in the whole-cell context can reveal morphological features of apposing organelles that suggest the effects of organelle packing.

Due to the apparent density of organelles in the cytoplasm and prevalence of organelle contacts, we hypothesize that organelles are subject to confinement forces constraining their morphology and position. A simple case of organelle packing occurs when one organelle expands, increasing the degree of organelle membrane packing within the cell and imposing size-dependent spatial constraints on all other organelles. Vacuole expansion is well documented in various cellular contexts, including in starvation, stress, cell cycle, drug treatment, osmotic stress, and cell stress or death (Aufschnaiter & Büttner, 2019; Rudge et al., 2004; Saravanan & Kane, 2025; Wang et al., 2001). It is a hallmark of some cancers, contributing to a “signet ring cell” phenotype in which an oversized vacuole fills the majority of the cytoplasm, while the remaining cellular contents are compacted at the periphery (Lossifides et al., 1980). In marine protists, vacuole inflation is a mechanism to maintain cellular buoyancy, illustrating that cellular vacuole content can fundamentally alter cellular physical properties (Larson et al., 2024). In budding yeast, vacuole enlargement mutants of the *vac, fab, atg,* and *vps* families were isolated in the late 1980s and 1990s (Banta et al., 1988; Yamamoto et al., 1995). The phenotype has been attributed to defects in vacuole PI(3,5)P2 regulation and fission, resulting in one fused vacuole, as well as increased vesicular trafficking to vacuoles in protein transport-defective mutants or upregulation of autophagy machinery (mTOR pathway activation) by metabolic stress, starvation or rapamycin (Bonangelino et al., 2002a; Dove et al., 2002; Jin et al., 2008; Rudge et al., 2004). Interestingly, it was previously noted that *fab1* temperature sensitive mutants exhibited morphological disruption of nuclear distribution and spindle morphology (Gary et al., 1998; Yamamoto et al., 1995).

The vacuole and nucleus are physically linked by the nucleus vacuole junction (NVJ), a membrane contact site consisting primarily of the Nvj1-Vac8 tethering complex (Kvam & Goldfarb, 2004; Pan et al., 2000). The NVJ is a metabolically responsive protein domain, with dynamic composition and size in response to cellular metabolic and stress conditions (Cadou & Mayer, 2015; Hariri et al., 2016, 2018; Tosal-Castano et al., 2021). It is a site of several metabolic processes, including lipid droplet biogenesis (Hariri et al., 2018), piecemeal microautophagy of the nucleus (Dawaliby & Mayer, 2010; Kvam & and Goldfarb, 2007; Kvam & Goldfarb, 2006), and lipid transfer at three-way junctions with mitochondria (Monteiro-Cardoso & Giordano, 2024). It is unclear whether all vacuole-nucleus interaction necessarily goes through tethering at NVJs, or whether NVJ formation can occur downstream of membrane-membrane proximity due to steric packing resulting from nucleus and vacuole size fluctuations.

The interplay between organelle size and intracellular geometry poses fundamental physical constraints that have only recently begun to be addressed through computational and theoretical frameworks. As organelle volumes scale with cell size, they necessarily reshape the spatial landscape that neighboring organelles occupy. Marshall (2020), in a theoretical treatment of subcellular scaling, demonstrated that growth of a large organelle such as the vacuole causes it to occupy a disproportionate fraction of cell volume, to the point of crowding neighboring organelles and displacing them from their resting positions (Marshall, 2020). These steric consequences of scaling are not merely positional, where physical contact between organelles also impose deformation forces that alter their shapes. At the molecular scale, Sui (2024) showed that excluded-volume interactions between polydisperse biomolecules, together with the spatial constraints imposed by the nuclear envelope, produce non-monotonic dynamics in the nuclear-to-cellular volume ratio under osmotic perturbation, providing a theoretical basis for understanding how steric forces propagate between the nucleus and the surrounding cytoplasm (Sui, 2024).

Together, these studies suggest that steric, confinement, and excluded-volume interactions are not incidental byproducts of organelle size but active physical determinants of organelle morphology and structural intracellular interactions.

In the present work, we focus on the mesoscale structural interaction between the vacuole and nucleus, the two largest organelles in budding yeast. We examine the aspects of the vacuole-nucleus relationship as pertains to their related morphological regulation at the whole-organelle/upper mesoscale: size regulation, shape, and interface geometries. We leverage our recently reported dataset of hundreds of high isotropic resolution single-cell reconstructions with multi-organelle information as a basis for statistically powerful analysis of morphological feature relationships between the vacuole and nucleus in a shared cellular context (Chen et al., 2025). This enables high-granularity 3D characterization of the effects of vacuole enlargement on nuclear morphology and cell anatomy overall.

Previous computational and theoretical works have treated the nucleus and cytoplasm as osmotically-nested compartments whose volumes are set by the balance of colloid osmotic pressures arising from differences in macromolecules concentration across the nuclear envelope (Deviri & Safran, 2022; Lemière et al., 2022a; Sui, 2024). While these models showed that nuclear size was governed by osmotic pressure differences, membrane mechanical properties, and the numbers of macromolecules that are too large to pass through the nucleus (Lemière et al., 2022a) and confine the nucleus (Sui, 2024), they lacked the spatial resolution necessary to capture the complex vacuole-nucleus morphological relationship within the cell. Recent work suggests a “nuclear drop model,” where the nucleus behaves like a liquid-filled drop bounded by a lamina that carries excess membrane area as folds and wrinkles(Dickinson, Abolghasemzade, and Lele, 2024); the model predicts two regimes: one where deformations cause unwrinkling, and another where no excess area remains and the nucleus is taut. Yeast nuclei, however, lack a lamina indicating that the nucleus morphology should instead be governed by the mechanical properties of the nuclear envelope (Meseroll and Cohen-Fix, 2016). To address questions pertaining to how confinement forces and inter-organelle packing shape the vacuole–nuclear morphological relationship, we employed a spatially-resolved modelling framework in which the nucleus, vacuole, and cell are each treated as osmotically active compartments, allowing us to obtain the morphological variation of the nucleus as a function of vacuole enlargement at the whole-cell level.

## Results

### Vacuole enlargement drives progressive disruptions in nuclear size scaling, positioning, and shape

It has been shown that cell shape variability can be a major factor complicating the morphometric modeling of organelles (42). We thus chose to use yeast due to their relatively simple and uniform cell shape, as compared to cultured animal cells. To further standardize organelle morphology analysis for statistical interpretability and modeling, we aimed to include only unbudded cells in our analysis. To enrich for unbudded cells in G1 phase, yeast cultures were grown to early log phase and rapidly plunge-frozen, preserving their ultrastructure. Cryopreserved cultures were then imaged by soft X-ray tomography in glass capillaries, creating 3D reconstructions at 30-35 nm isotropic resolution.

Our core dataset of control and large-vacuole yeast imaged with soft X-ray tomography and automatically segmented has been described in depth in our previous work (39) but is briefly described here. Three closely related *Saccharomyces cerevisiae* strains were used: the haploid wild-type strain BY4741A, a version of the parental strain expressing the vacuole membrane marker VPH1-GFP, and a large vacuole mutant, *vac14* (16,22), also expressing VPH1-GFP. From these datasets, single cells of suitable quality and cell cycle phase were selected, and cell, vacuole, nucleus, and lipid droplet compartments were auto-segmented based on a custom CNN. Following quality checking, our curated dataset contains 287 WT cells, 138 VPH1-GFP cells, and 65 *vac14* cells, a total of 490 cells.

We have previously demonstrated that *vac14* exhibits a general vacuole size variability phenotype with an enrichment of large-vacuole cells, also driving global enlargement of cells and nuclei (Chen et al., 2025). In other words, many *vac14* cells exhibit normal vacuole morphology at G1, with the phenotype generally progressing with cell growth. Interestingly, this mutant displays normal viability and growth at 30 °C despite the vacuole enlargement phenotype (Bonangelino et al., 2002b), while mutations in functionally related genes, such as FAB1, an upstream regulator of VAC14, trigger a stronger vacuole enlargement phenotype that is lethal and necessitates temperature-sensitive mutants for study (Gary et al., 1998). Additionally, vacuole size variability in other contexts can be a feature of normal homeostasis, such as in normal cell growth and division (Matos-Perdomo et al., 2022; Uchida et al., 2011), non-lethal cellular stress response, such as starvation or osmotic shock (Bonangelino et al., 2002b; Saravanan & Kane, 2025), or of significant dysfunction or disease states, such as cancer or cell death (Lossifides et al., 1980; Mao et al., 2025; Wen & Ma, 2025). This illustrates that there is a range of vacuole size that is well-tolerated by cells, above which basic cellular function is impaired. We therefore hypothesize that below this viability threshold, vacuole size variability encompassing normal ranges and *vac14-*triggered enlargement is accompanied by a range of morphological effects on other organelles such at the nucleus, and that quantifying the magnitude of these effects can reveal the distinct physical constraints on specific nuclear features, such as size, position, and shape. Since this hypothesis is based on vacuole size and not genotype, we enriched for enlarged vacuoles in our dataset by pooling all 490 cells and dividing the dataset by “normal” and “high” vacuole:cell volume ratio (which we will denote as V:C). The threshold for this cutoff was defined as 1 standard deviation above the mean for V:C (0.27, based on mean 0.18 and standard deviation 0.9). This produced our comparison datasets which we will refer to as normal V:C (<0.27) and high V:C (>0.27).

Voxel-based per-cell volume measurements for each structure are shown for the full pooled dataset, normal V:C, and high V:C cells, demonstrating a strong separation in volumetric phenotype between the normal and high V:C groups (Fig. S1, Table S1). Most cells in our dataset exhibit one vacuole in log phase, with no more than 5 distinct vacuoles observed in a few instances, enabling straightforward morphological analysis of the vacuole-nucleus relationship. Vacuole volume analyses are reported as the total per-cell volume (“Vacuole Sum Volume”). Slicing the dataset by V:C demonstrates the portion of the total distribution covered by the smaller, high V:C group. Notably, all compartments show a general increase in volume in the high V:C group. In particular, the long right tail in cell and vacuole whole-dataset distributions is entirely attributed to the high V:C group (Fig. S1A, B). A smaller but significant shift is seen in the nucleus and LD volume distributions (Fig. S1C, D). The proportional volume composition comparison for each group shows that the vacuole occupies 18-25% of cell volume in pooled (“All”) and normal (“V:C <0.27”) groups, and over 35% in large-vacuole (“V:C > 0.27”) groups.

We first asked how vacuole and nucleus size scaling behaviors compare between normal and high V:C groups. While our data are grouped by vacuole:*cell* (V:C) volume ratio to reflect a conventional proportional relationship to overall cell size, our scaling analysis focuses on the relationship between organelle and *cytoplasmic* volumes, where the cytoplasm is defined as the cellular volume minus the volume of the organelle in question (vacuole or nucleus). This is to remove the confounding self-correlation of the organelle of interest as part of the total cell volume, enabling us to directly assess each organelle’s relationship with the rest of the cell. The cytoplasmic compartment devoid of known organelles, which we refer to as “free cytoplasm”, “free space”, or “Cell-Organelles”, can be considered a proxy for crowdedness, with the caveat that this space contains many other organelles and cellular components of unknown size and distribution.

Log-log scaling plots demonstrate distinct scaling trends for each compartment in normal and high V:C conditions. For normal V:C cells, vacuole volume displays supralinear scaling (alpha 1.84) and nucleus volume displays nearly linear scaling (alpha 0.86) with cytoplasmic volume (cell volume excluding the vacuole or nucleus, respectively). This is consistent with previous reports for the yeast vacuole (Chan et al., 2016; Chan & Marshall, 2014) and nucleus (Gillooly et al., 2015; Jorgensen et al., 2007), but unlike previously published work, demonstrates both phenomena in the same dataset, with the caveat that the choice of parental strain and restriction to early G1 cells in our dataset may introduce small differences between our and previous reports. In normal and high V:C cells, the vacuole scaling factor is consistent (normal V:C 1.84, high V:C 1.79), but the correlation is stronger in high V:C (normal r^2^ = 0.64, high r^2^ = 0.76), suggesting that vacuole and cytoplasm volume are more tightly linked in cells with enlarged vacuoles. Conversely, the nuclear scaling factor and correlation coefficient are reduced in high V:C cells (normal alpha 0.86, r^2^ = 0.53, high V:C alpha 0.53, r^2^ = 0.26), demonstrating that vacuole enlargement breaks nuclear size scaling. Free cytoplasm scales strongly positively with organelles (Fig. S2A), but proportionally scales negatively with cell volume, particularly in high V:C (Fig. S2B). Since vacuole enlargement is associated with an increase in cell size, we interpret these results as showing that nuclei and free cytoplasm grow with cell size but at a reduced rate rather than decreasing in size.

Do the proportional occupancy of vacuoles and nuclei remain constant or change in large-vacuole cells? To directly visualize the effect of vacuole enlargement on organelle volumetric ratio, we plot the ratio of vacuole and nucleus size to cytoplasmic volume, i.e. the part of overall cell volume that is left after the organelle is removed (denoted by Vac:Cyto and Nuc:Cyto respectively) against the cytoplasmic volume, including (Figure 1D, E) or excluding the organelle of interest (Fig. S2C, free space). In each case, we plot the data after separating into groups with normal and high V:C ratios (which reflect the more standard way of defining scaling relative to overall cell volume). Vacuole to cytoplasm ratio increases steadily with cytoplasmic volume in both normal and high V:C, with a stronger association in high V:C (r^2^ = 0.38 vs 0.27). Nucleus volume to cytoplasm ratio is roughly constant with cytoplasmic volume in normal V:C, but becomes negatively correlated with cytoplasmic volume in high V:C cells.

**Figure 1.**
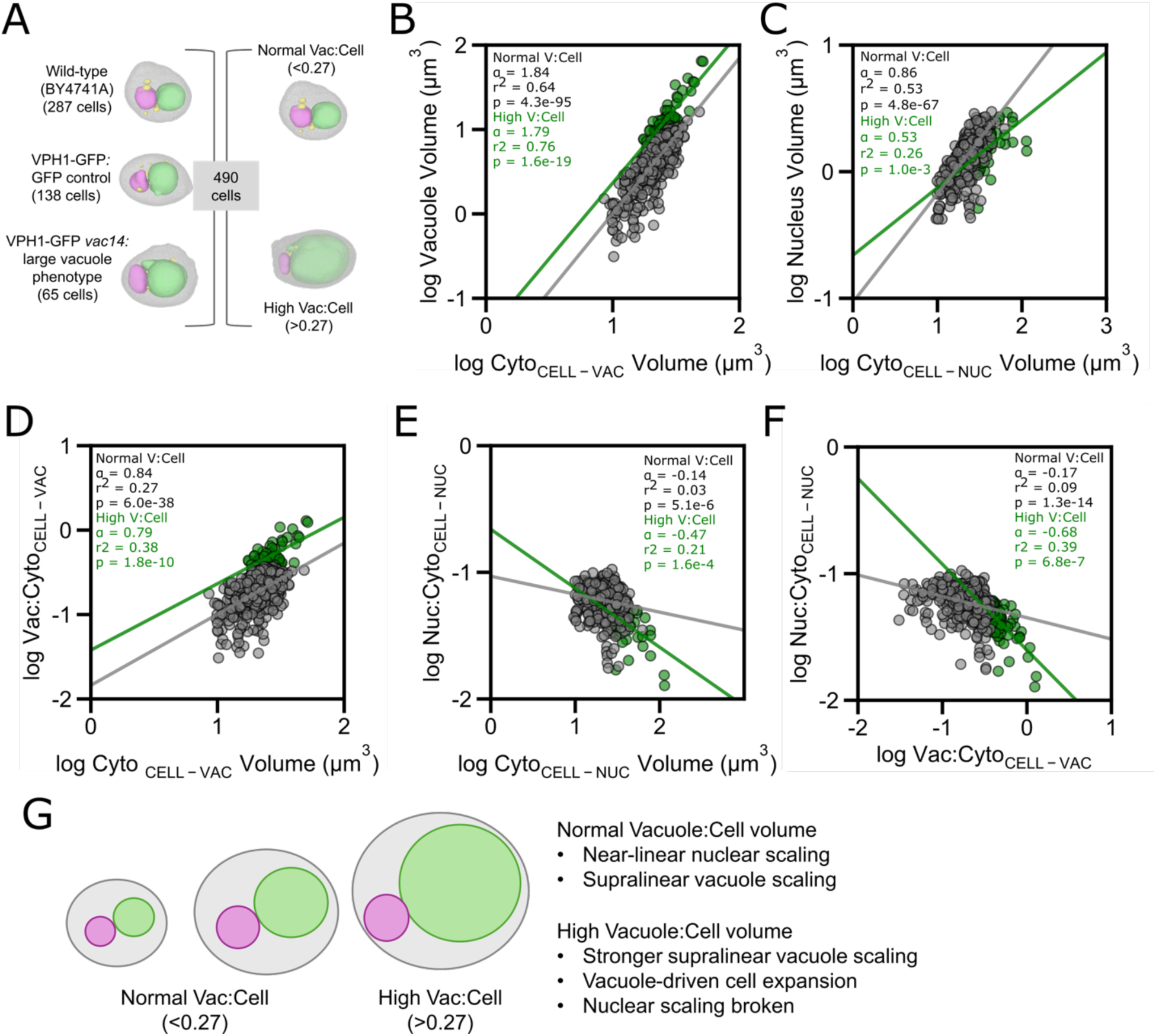
Vacuole enlargement breaks nuclear size scaling. **A)** The dataset consists of whole-cell 3D reconstructions of soft X-ray tomograms of individual unbudded yeast cells from 3 strains: wild-type parental strain BY4741A, VPH1-GFP (parental with GFP tag), and *vac14* (knockout strain containing VPH1-GFP marker). Images shown here are triangulated mesh renderings of individual cells including cell (grey), vacuole (green), nucleus (pink), and lipid droplet (yellow) segmentations. **B-F)** Log-log scatterplots of inter-compartmental volume scaling relationships. **B)** Vacuole volume scaling with cytoplasm, where cytoplasm is defined as cell volume excluding vacuole volume. **C)** Nucleus volume scaling with cytoplasm, where cytoplasm is defined as cell volume excluding nucleus volume. **D)** Vacuole:Cytoplasm (Cell-Vacuole) scaling with cytoplasm (Cell-Vacuole). **E)** Nucleus:Cytoplasm (Cell-Nucleus) scaling with cytoplasm (Cell-Nucleus). **F)** Nucleus:Cytoplasm (Cell-Nucleus) scaling with Vacuole:Cytoplasm (Cell-Vacuole). **G)** Schematic summarizing scaling phenomena in normal and enlarged vacuole (high V:C) contexts.

We also looked at whether there is a direct scaling relationship between vacuoles and nuclei (Fig.1F). In normal V:C cells, there is no association between nuclear and vacuolar volume ratios, but in high V:C cells, a moderate negative association is observed, suggesting that vacuole size above the normal threshold impinges on nuclear scaling. This is also visible in bulk statistics of proportional volume composition (Fig. S1E). Therefore, as vacuoles occupy a larger proportion of cytoplasmic space, nuclei occupy a smaller proportion.

In addition to altered size scaling, another potential effect of vacuole enlargement could be spatial displacement of the nucleus. Nuclear shift toward the cell periphery accompanying vacuole enlargement has been anecdotally reported in a variety of contexts, including budding yeast *fab1* mutants (Gary et al., 1998), dinoflagellates (Larson et al., 2024), and follicular lymphoma. We asked whether nucleus positioning relative to the cell geometric center was dependent on vacuole size in our dataset. Nuclear position was calculated as the Euclidean distance between nuclear and cell 3D centroids and normalized by cell Feret diameter. A plot of the distance between nucleus centroid and cell centroid showed a general increase in nucleus-cell centroid distance with V:C (Fig. 2). Interestingly, we find a moderate association between nuclear position and V:Cyto (Fig. 2B) or proportional free cytoplasm (Cell-Organelles:Cell, Fig. 2C) in normal but not large-vacuole cells. These results suggest that the nucleus is positioned further from the cell center depending on vacuole size or available space even in morphologically normal cells, but vacuole enlargement may limit the space available for the nucleus to occupy as it approaches the cell boundary.

**Figure 2.**
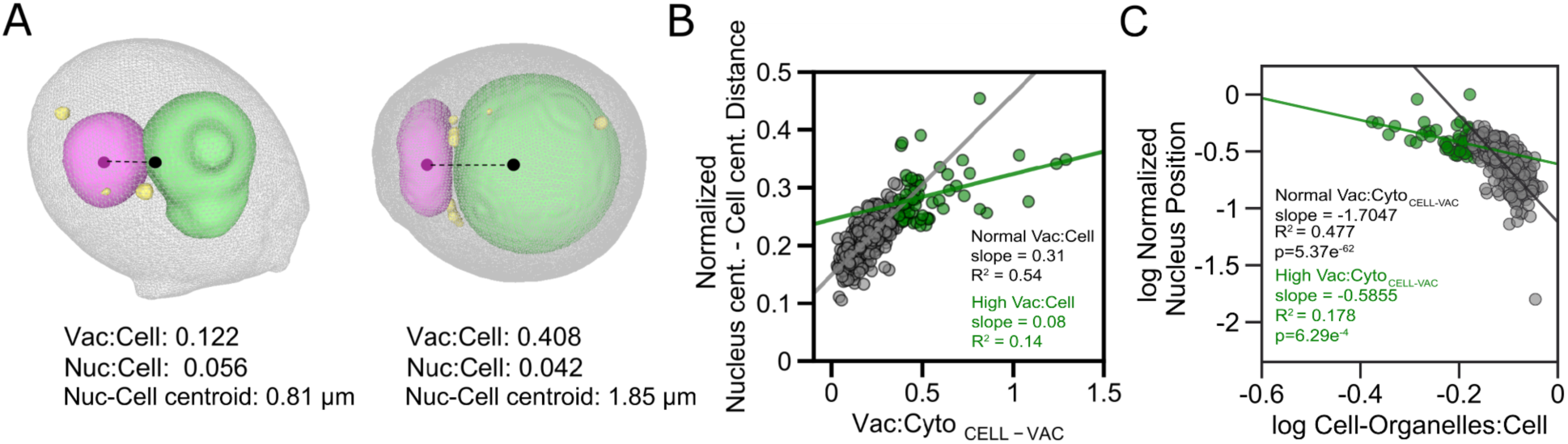
Vacuole enlargement displaces the nucleus. **A)** Representative mesh-rendered cells showing nuclear positioning in a cell with a normal vacuole (left) and enlarged vacuole (right). Grey - cell, green - vacuole, pink - nucleus, and yellow - lipid droplet. **B, C)** Log-log scatterplot of correlation between nuclear position (distance between nuclear and cell centroids, normalized by cell Feret diameter) and **B)** V:Cyto ratio for normal (black/grey) and high (green) V:C cells and **C)** Cytoplasm:Cell ratio.

A particularly striking observation in very large vacuole cells is the apparent deformation of the nucleus, seen in our data (Fig. 3A), as well as elsewhere (Bianco et al., 2023; Gary et al., 1998). We hypothesized that if the vacuole were directly responsible for nuclear shape change due to spatial packing, we should see a progressive flattening of the nucleus with vacuole enlargement. In bulk, nuclear sphericity is significantly lower in normal versus high V:C cells, reflecting nuclear flattening (p=5.95e^-8^). The vacuole also showed a slight distribution shift toward more spherical vacuoles in high V:C cells, likely reflecting fusion as well as expansion of vacuoles, although the overall difference in sphericity is not significant (p=0.3) (Fig. 3B). This led us to hypothesize that the vacuole and nucleus may engage in “shape competition” when packing is in effect, such that one, the nucleus, deforms away from a sphere as the vacuole becomes both larger and more spherical.

**Figure 3.**
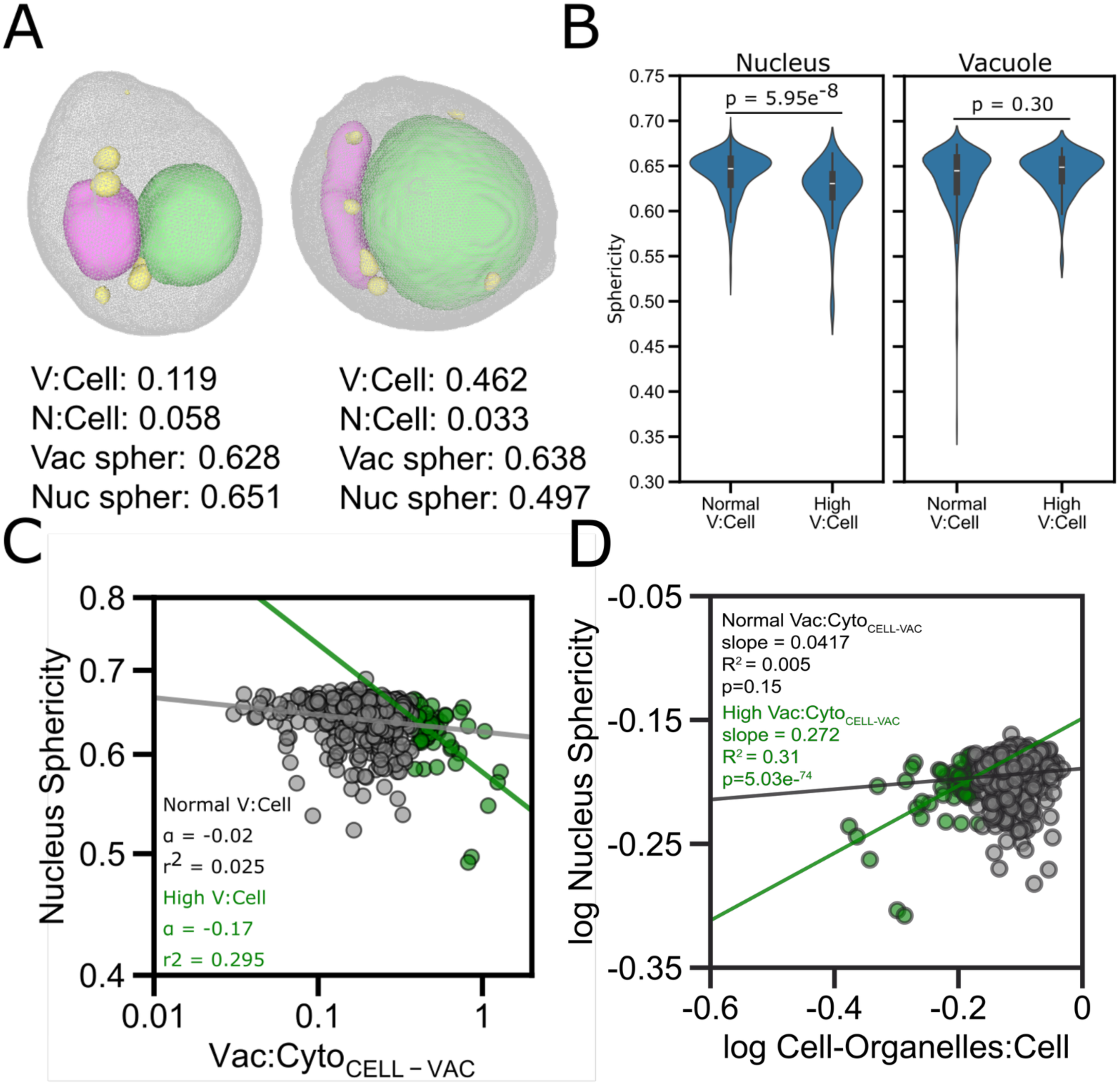
Nuclear shape is associated with V:Cyto volume ratio, dependent on vacuole enlargement. **A)** Representative mesh-rendered cells showing nuclear positioning in a cell with a normal vacuole (left) and enlarged vacuole (right). Grey - cell, green - vacuole, pink - nucleus, and yellow - lipid droplet. **B)** Violin plots of voxel-based sphericity distributions for nucleus and vacuole in normal and high V:C groups. **C, D)** Log-log scatterplot of correlation between nuclear sphericity and V:Cyto ratio for normal (black/grey) and high (green) V:C cells.

Our shape competition hypothesis predicts that the ability of one organelle to deform another would be greatest when there was less available space inside the cell. Nuclear sphericity had no correlation with either V:Cyto or available cytoplasmic space (Cell-Organelles:Cell, cell volume excluding known vacuole and nuclear volumes) in normal V:C cells (Fig. 3C-D). In high V:C cells, however, nuclear sphericity was negatively correlated with V:C and positively correlated with cytoplasmic space. This suggests that nuclear shape is constrained by free space, which becomes limited by vacuole size in high V:C space, in other words demonstrating vacuole-driven nuclear deformation which only occurs when the vacuole occupies an abnormally high proportion of the cell.

### Simulations recapitulate effects of vacuole enlargement on nucleus

We have shown that vacuole enlargement is associated with various morphological effects on the nucleus, including dysregulating size scaling (Fig. 1), positioning (Fig. 2), and shape (Fig. 3). How do these effects relate to each other? Is there a vacuole size-dependent order to the onset of each effect? Pinpointing a precise relative relationship between these phenotypes, such as a specific vacuole size threshold associated with each nuclear deviation from normal scaling, could reflect varying degrees of robustness of each nuclear property to vacuole expansion and suggest a physical mechanism underlying vacuole-nucleus morphological interaction. Figure 4A shows a selection of individual cells arranged by increasing V:C volume ratio. Table 1 displays the values for V:C, N:C, vacuole and nuclear sphericity, and nuclear position for each cell. The values are marked in bold where they fall outside the normal range, defined as 1 standard deviation above or below the mean.

**Figure 4.**
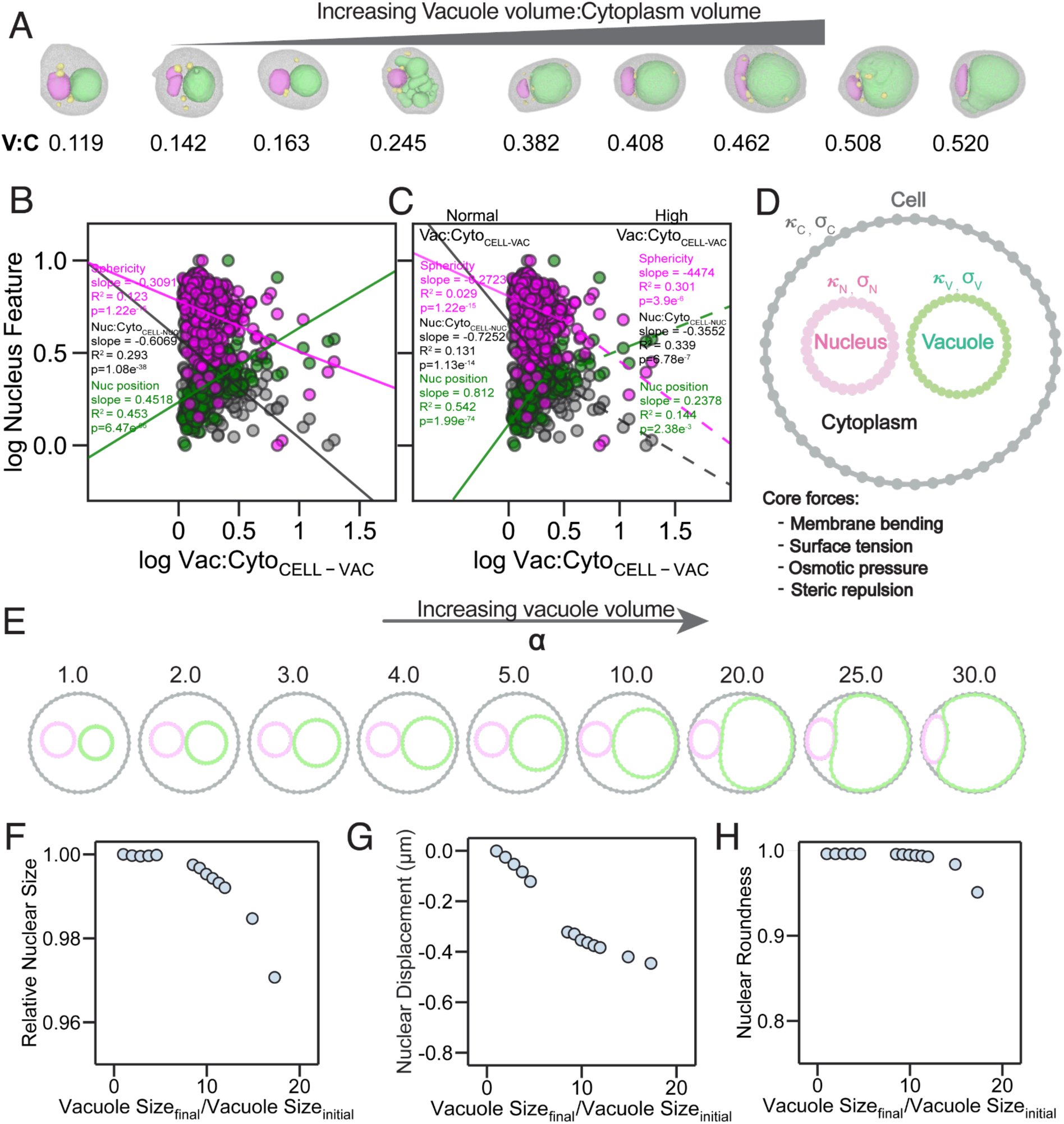
Vacuole size-dependent effects on the nucleus can be recapitulated by a biophysical simulation. **A)** Gallery of single cell mesh reconstructions selected to illustrate progressive onset of nuclear morphological effects associated with increasing V:C volume (left to right). Gray - cell, green - vacuole, pink - nucleus, yellow - LDs. Additional metrics in Table 1. **B)** log-log scatterplots showing correlations of the nuclear sphericity (magenta), Nuc:Cyto volume ratio (grey), and nuclear position (green) to V:Cyto volume ratio. **C)** log-log scatterplots showing correlations of the nuclear sphericity (magenta), Nuc:Cyto volume ratio (grey), and nuclear position (green) to normal and high V:Cyto volume ratio **D)** Schematic representation of the biophysical model showing the nucleus in pink, vacuole in green and the cell in grey. Cell and organelles are modeled as discrete 2-D polygons with bending rigidity () and surface tension (). Equilibrium size of the organelle (or cell) is determined by balance between mechanical properties and the osmotic pressure of the organelle (cell). **E)** Snapshots showing the final equilibrium morphologies of the nucleus, vacuole and cell after minimization using conjugate gradient varying the vacuole enlargement (left to right) and increasing the vacuole bending rigidity relative to nucleus bending rigidity (top to bottom). **F)** Nuclear size relative to the initial size, **G)** nuclear roundness , and **H)** nuclear displacement relative to cell centroid as a function of the vacuole enlargement constrained by osmotic and other forces.

**Table 1.** Metrics underlying cells displayed in Figure 4A. Normal range (+/- 1 standard deviation from mean) for each metric is indicated in parentheses in the first column. Values falling outside this range are indicated in bold.

| Metric | Cell 1 | Cell 2 | Cell 3 | Cell 4 | Cell 5 | Cell 6 | Cell 7 | Cell 8 | Cell 9 |
| --- | --- | --- | --- | --- | --- | --- | --- | --- | --- |
| Vacuole:Cell<br>(0.103-0.269) | 0.119 | 0.142 | 0.163 | 0.245 | **0.382** | **0.408** | **0.462** | **0.508** | **0.520** |
| Nucleus:Cell<br>(0.041-0.070) | 0.058 | **0.019** | 0.067 | **0.040** | **0.020** | 0.042 | **0.033** | **0.021** | **0.025** |
| Vac Sphericity<br>(0.603-0.670) | 0.646 | 0.647 | 0.666 | **0.362** | 0.649 | 0.654 | 0.638 | 0.651 | **0.546** |
| Nuc Sphericity<br>(0.611-0.665) | 0.651 | **0.522** | 0.689 | 0.651 | 0.616 | **0.59** | **0.497** | 0.626 | **0.546** |
| N-C normalized<br>distance ($\mu\text{m}$)<br>(0.178-0.257) | **0.167** | 0.182 | 0.193 | 0.218 | 0.237 | **0.288** | 0.256 | **0.308** | **0.276** |

When we visualize the variation of all three nuclear features (volume ratio, position, and sphericity) with V:C simultaneously by normalizing their values to observed ranges, the relative slopes indicate a distinct order of onset of nuclear morphology phenotypes with vacuole enlargement. Among the entire pooled dataset (Fig. 4B), nuclear volume ratio declines fastest (slope = -0.61), followed by nuclear displacement (increase in distance from cell center, slope = 0.45), and a more gradual decrease in sphericity (slope = -0.31). The correlation coefficients are low to moderate for all three with high statistical significance (r^2^ = 0.293 (Nuc:Cyto ), 0.123 (sphericity), 0.453 (position). When separating regression lines by the high V:C threshold, nuclear volume ratio and position are more correlated with V:Cyto in normal V:C cells than in high V:C cells. In contrast, sphericity is uncorrelated with V:Cyto in normal V:C cells (slope = - 0.27, r^2^ = 0.03), but is correlated in the high V:C subset (slope = 0.45, r^2^ = 0.30) (Fig. 4C).

To further investigate the physical mechanisms underlying how vacuole enlargement and space constraints might affect the size scaling, positioning, and shape effects of the nucleus, we employed computer simulations to study the morphologies of the nucleus as a function of vacuole size. Inspired by work of Lemiere et al. (2022) and experimental observations that expansion of the vacuole can be driven by the high osmotic pressures due to calcium transport within the vacuole (Baars et al., 2007; Chan & Marshall, 2014; Wilson et al., 2018), we modeled the organelles and cell as enclosed osmotic compartments which can swell and shrink. Here, we incorporated established theoretical work in a spatially resolved mechanical framework to capture the morphology of the organelles. Briefly, we modeled the organelles as discrete 2-D enclosed polygons with resistance to bending and stretching, and osmotic pressure (Fig. 4D, see Methods for more details) (Nakamura et al., 2023; Lee and Rangamani, 2024; Zhu et al., 2022). To examine the impact of organelle packing, we account for excluded volume interactions between organelles and the cell using a repulsive potential. To systematically vary the size of the organelles, we modulated the osmotic pressure of each organelle using the established van’t Hoff relationship (Hoff, 1887). Here the preferred organelle size is altered by changing the solute concentration within the organelle. We initialized the nucleus, vacuole and the cell as ellipses. The vacuole and nucleus are confined within the cell and not in direct contact (Fig. 4D).

Since our computational model operates in two dimensions, we define a scaled volume, 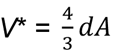, to convert between three-dimensional volumes and our two-dimensional area in a manner that preserves the geometric scaling of each compartment. Here 𝑑 is the average distance from the centroid to the edge of the 2-D polygon. This mapping assumes spherical compartment geometry, such that the scaled volume corresponds to the cross-sectional area of a sphere whose volume equals the measured three-dimensional volume. The initial organelle scaled volumes are derived from the experimentally reported values of organelle volumes. The parameters for the model were derived from the existing literature (cf. Table 2). To modulate the target size of the vacuole, we systematically varied the target scaled vacuole volume as 𝛼V, where 𝛼 ranges from 1 to 30, and V is the initial vacuole volume. We evolve this system towards mechanical equilibrium using the conjugate gradient method. Further analysis on the nuclear morphology was performed using the equilibrium organelle morphologies. Given the symmetry of the experimental system, a 2-D cross-section of the 3-D cell captures the essential geometry of the system. Hence, we can take advantage of this symmetry to capture the relevant physics and morphological information of interest without the computational overhead of 3-D simulations.

**Table 2.** List of parameters used in simulations.

| Parameter (units) | Value | Reference |
| --- | --- | --- |
| Cell initial volume, $V_C$ ($\mu\text{m}^3$) | 24.22 | Parametrized for this study |
| Nucleus initial volume, $V_N$ ($\mu\text{m}^3$) | 0.69 | Parametrized for this study |
| Vacuole initial volume, $V_V$ ($\mu\text{m}^3$) | 0.69 | Parametrized for this study |
| Cell $\underline{V}$ | $V_C$ | Parametrized for this study |
| Nucleus $\underline{V}$ | $V_N$ | Parametrized for this study |
| Vacuole $\underline{V}$ | [1, 2, 3, 4, 5, 10, 11, 12, 13, 14, 15, 20, | Varied for this study |
| | $25, 30] \times V_v$ | |
| Cell tension, $\sigma_C$ (N/m) | 10 | Lemiere et al. 2021 (Lemière et al., 2022b) |
| Nucleus tension, $\sigma_N$ (mN/m) | 0.001 | Lemiere et al. 2021 (Lemière et al., 2022b) |
| Vacuole tension, $\sigma_V$ (mN/m) | 0.001 | Lemiere et al. 2021 (Lemière et al., 2022b) |
| Cell bending rigidity, $\kappa_C$ ($k_B T$) | 1000 | Parametrized for this study |
| Nucleus bending rigidity, $\kappa_N$ ($k_B T$) | 30 | Lim et al. 2007 (Lim H. W. et al., 2007) |
| Vacuole bending rigidity, $\kappa_V$ ($k_B T$) | $[1, 10, 50, 100] \times \kappa_N$ | Varied for this study |
| Cell osmotic strength constant ($nN\mu m$) | 1.0 | Parametrized for this study |
| Nucleus osmotic strength constant ($nN\mu m$) | 0.1 | Parametrized for this study |
| Vacuole osmotic strength constant ($nN\mu m$) | 0.1 | Parametrized for this study |
| Repulsive potential strength, $K^{rep}$ ($pN\mu m$) | $4 \times 10^{-8}$ | Floyd et al. 2021 (Floyd et al., 2021) |
| Tether spring constant ($pN/\mu m$) | 0.1 | Parameterized for this study |
| Tether equilibrium length ($\mu m$) | 0.03 | Eisenberg-Bord et al., 2016; Hariri et al., 2018 |

Figure 4E shows snapshots of the equilibrium morphologies of nucleus, vacuole and cell wall from simulations. 𝛼 = 1 serves as the baseline in which vacuole size and nuclear shape are unchanged. With a modest increasing 𝛼 < 10, we observed larger vacuoles which displaced the nucleus towards the cell wall, however without pronounced nuclear deformations by visual inspection. When 𝛼 > 10, we observed the onset of nuclear deformation. Our simulations indicated that nuclear deformation occurred when the combined vacuole and nucleus size was sufficiently large to contact both sides of the cell wall, suggesting that spatial constraints induced by vacuole enlargement plays a role in nuclear morphological deformations.

To quantify the impact of vacuole enlargement on nuclear morphology, we calculated the nuclear size, displacement relative to cell centroid, and roundness as a function of vacuole size. We calculated the nuclear size as the nuclear area divided by the initial area (Fig. 4F). Values smaller than 1 indicate smaller nuclei. Next, we examined how vacuole enlargement impacted displacement of the nucleus (Fig. 4G). We calculated the nuclear center of mass (c.o.m.) displacement relative to the displacement of the cell center of mass. Negative values in the nuclear displacement indicate a leftward shift in the nucleus relative to the cell. Lastly, we use roundness as a proxy measure for sphericity in 2-D to capture deviations from a perfect circle, where values closer to 1 indicate a perfect circle and values smaller than 1 indicate deviation from a circle. We computed the nuclear roundness as the nuclear perimeter divided by the nuclear area (Fig. 4H).

Initially, when the vacuole enlargement is small, nuclear size and roundness are uncorrelated with vacuole size, while nuclear displacement decreases. As the vacuole becomes larger, nuclear size decreases in agreement with experimental findings. There is an accompanying leftward shift in the c.o.m. of the nucleus relative to the cell indicating that the nucleus is decentralized. Our simulations also suggest that different extents of vacuole enlargement are required for the onset of different nuclear morphological deviations. Decreases in nuclear size and displacement begin to occur at smaller vacuole enlargement (Figs. 4F and G) . In contrast, nuclear roundness remains largely unchanged at modest vacuole enlargements, only decreasing at much larger enlargements (Fig. 4H). These results recapitulate the experimental nuclear deformation phenotype and demonstrate that vacuole-driven spatial confinement is sufficient to produce the observed shifts in nuclear size and displacement (cf. Fig. 4C).

### Integrated shape modeling represents morphological interplay between vacuole-driven nuclear defects and reveals covarying feature sets

Our analyses so far have focused on the relationship between pre-defined, hypothesis-driven organelle parameters: vacuole volume ratio, nucleus volume ratio, nucleus position, and nucleus sphericity. However, this type of analysis leaves the possibility open that there may be other meaningful features that more directly reflect the relationship between nucleus and vacuole morphologies. For a less biased approach, we adapted a joint integrated shape modeling methodology developed by the Allen Institute for Cell Science (Viana et al., 2023) to approximate the shapes of the cell, vacuole, and nucleus for each cell using spherical harmonics expansion. Single cell three-channel binary z-stacks were aligned with the nucleus-vacuole centroid axis along the horizontal plane and the nucleus centroid as the image center. The cell, vacuole, and nucleus shapes are then modeled using a spherical harmonic expansion, identifying a set of coefficient values for the spherical harmonic terms that together can recapitulate the simplified 3D shape of the structure without losing key features. This produces a set of 289 spherical harmonics coefficients for each structure reflecting its deviation from a sphere. To examine the co-variation of all shape descriptors for each cell, the coefficients are pooled, resulting in 867 dimensions per cell. Principal component analysis of these coefficients for experimental images of all cells was used to produce a 5-dimensional shape space capable of distinguishing the top 5 shape modes explaining the majority of observed variance. The center of the 5D space is represented by a generated cell-vacuole-nucleus reconstruction that can be considered the average configuration, generating an ellipsoidal cell shape, with a large spherical vacuole in close proximity to a smaller spherical nucleus, situated at the center of the cell. Sampling at various points on each principal component axis reveals a range of cell configurations varying around this average, enabling human-interpretable visualization of co-varying features and how they contribute to cell-to-cell morphological variance across the dataset (Fig. 5A).

**Figure 5.**
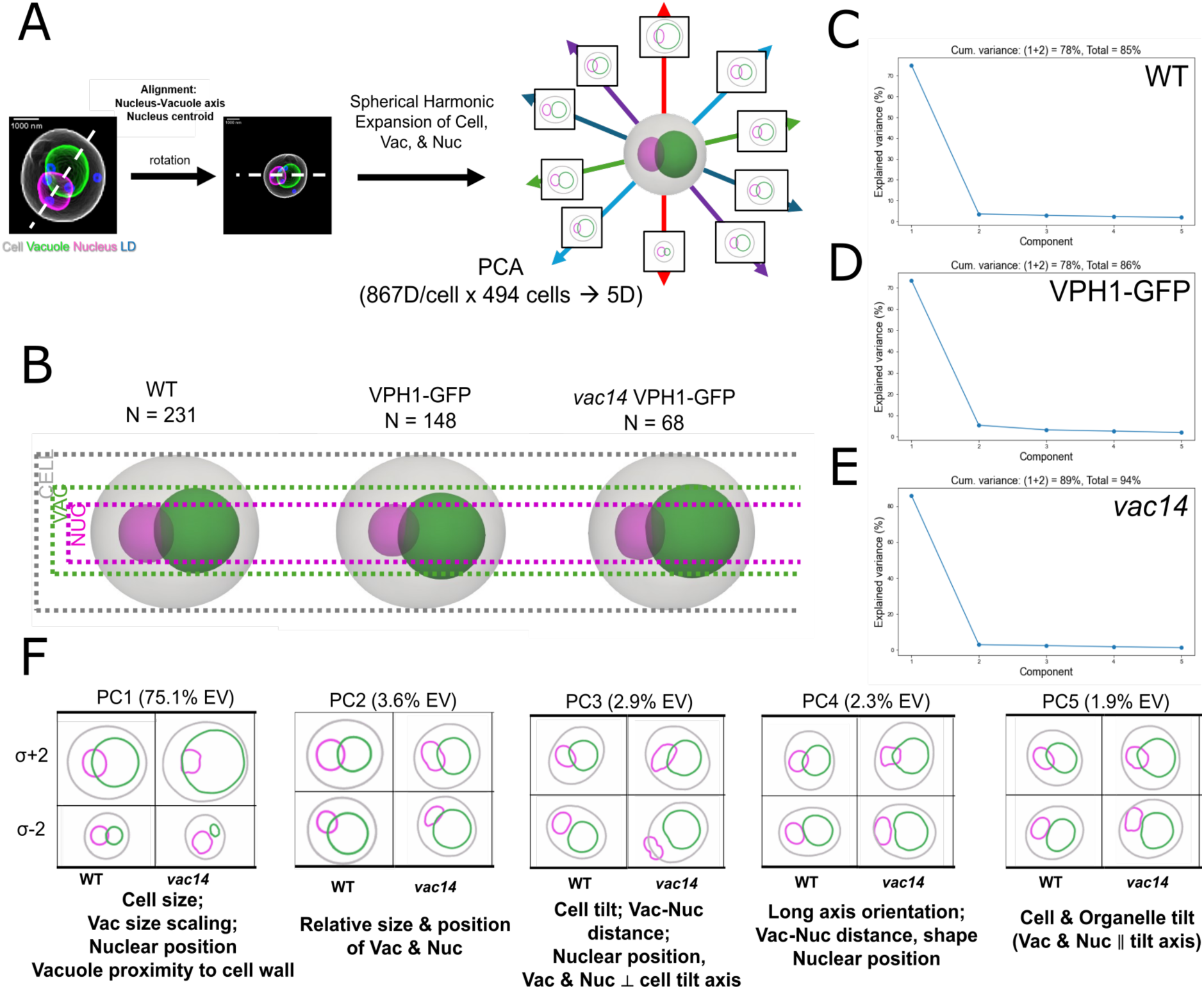
Integrated shape modeling of cell/vacuole/nucleus morphological co-variation demonstrates spatial confinement as a driving factor of vacuole-nucleus interaction. **A)** Schematic illustrating modeling pipeline. Single-cell multi-channel 3D images containing segmentations for cell, vacuole, and nucleus are cropped and rotated to align along the nucleus-vacuole centroid axis. Spherical harmonic expansion is applied to each cell to represent each structure by 289 coefficients, producing a 867-dimensional set of coefficients per cell. Principal component analysis (PCA) projects the dataset onto a 5-dimensional shape space which represents >=85% of morphological covariation. Theoretical cell configuration instances can be generated throughout the shape space, with the origin representing an “average cell”. **B)** Average cell configurations for integrated shape models generated for each strain, visually normalized by cell size to show subtle variations in nuclear and vacuole size. **C-E)** Scree plots showing the cumulative variance and relative contribution of each of the 5 principal components isolated for each strain model. **F)** Side-by-side comparison of principal components for WT (Supplementary Video 1) and *vac14* (Supplementary Video 2) models. Top and bottom rows represent generated images at 2 standard deviations above and below the mean respectively, providing opposing representations as a visual identification of co-varying morphological features.

The workflow can be applied to subsets of the dataset to examine differences in morphological trends between sub-populations of cells, provided the subpopulation is of sufficient size and variability to populate an even distribution of bins. An average cell configuration is shown for each strain within our dataset (Fig. 5B). Overall, the average configurations reflect only subtle differences between the strains when visualized side by side with normalized cell size. The nucleus is slightly smaller and shifted relative to the vacuole in VPH1-GFP as compared to wild-type and *vac14*, and the *vac14* nucleus is slightly larger than both strains. This is consistent with our previous observations that the VPH1-GFP strain has slight morphological differences to WT, despite the presumed lack of functional effect of VPH1-GFP transgene expression, whereas *vac14* deletion in a VPH1-GFP background produces a vacuole size variability phenotype with only subtle effect on average vacuole size (Chen et al., 2025) .

For each strain, the first principal component accounts for over 75% of explained variance, with the other four components explaining 1-3% of variance (Fig. 5C-E). Well populated, Gaussian-like histograms and low Spearman correlation coefficients between all principal components indicate that the components are well-separated, reflecting distinct feature combinations in the morphospace. We note that the distribution is noisier and correlation coefficients elevated slightly higher in *vac14*, reflecting the lower sample size in this subset.

2D visualization of cell-vacuole-nucleus configurations between -2 and +2 standard deviations from the mean allow us to interpret the morphological features co-varying within each shape mode (Fig. 5F, Supplementary Videos 1 and 2). PC1, which accounts for an overwhelming amount of explained variance (EV) (75.1% WT, 86.0% *vac14*), is primarily marked by size scaling of vacuole and nuclei in all strains. The size differential between nucleus and vacuole in the WT +2 standard deviation image reflects the differential organelle scaling rates. The difference is more pronounced in *vac14*, reflecting the vacuole enlargement phenotype that arises in a subset of these cells. Interestingly, we note a slight irregularity in nuclear shape in the *vac14* image, but not in WT. These results confirm the expected result that organelle size scaling plays a dominant role in determining morphological relationships and cell anatomy patterns when cell shape is constrained, as previously shown in hiPSCs (Viana et al., 2023), which have highly variable cell shape compared to cell walled yeast cells. At the same time, they also reflect more subtle features that correlate with scaling behavior, such as vacuole membrane proximity to the cell periphery and relative positioning of organelles.

The remaining PCs show the coupling of features other than organelle-cell size scaling. PC2 (EV - 3.6% WT, 3.0% *vac14*) reveals covariation between the relative organelle sizes and their spatial arrangement. At σ+2, vacuole and nucleus are nearly equivalent in size and oriented parallel, but at σ-2, the vacuole is larger than the nucleus, and the organelles are positioned at an angle relative to the long axis of the cell. In *vac14*, this trend is accompanied by morphological variation in the nucleus, which appears symmetrically flattened at σ+2, but flattened specifically at the vacuole interface at σ-2. PC3 (EV - 2.9% WT, 2.4% *vac14*) reveals covariation between inter-organelle spacing, cell tilt, organelle tilt, and organelle shape. The cell appears more elongated than in PC1 and PC2. Given the rigidity of the yeast cell wall, we speculate this mode might reflect an influence of cell shape on organelle geometry. At σ+2, the vacuole-nucleus axis is nearly perpendicular to the cell long axis, the organelles are tightly in contact. Interestingly, organelles appear slightly flattened at the vacuole-nucleus interface in WT, but in *vac14* the vacuole appears to retain a round shape, while the nucleus is flattened at the interface. At σ-2, the organelles are arranged parallel and spaced apart. Here, a slight flattening at the interface is seen in both organelles in both WT and *vac14*, with more significant deformation in the nucleus in *vac14*. In both, the organelles themselves have a tilted appearance, aligned with the cell’s long axis, which shifts between sigma +2 and -2. PC4 (EV - 2.3% WT, 1.8% *vac14*) and PC5 (EV - 1.9% WT, 1.3% *vac14*), show similar relationships between organelle shape and orientation with respect to each other and the cell long axis. Results for the VPH1-GFP strain are very similar to WT (Supplementary Video 3).

Interpreting these configurations requires caution, given that they are generated synthetic images of points derived from linear trends in a shape space, rather than representations of real cells. Therefore, some generated configurations may not exist in the dataset, but sit on a principal component axis opposite realistic morphologies, such as the tiny vacuole seen in *vac14* PC1, σ-2, and the apparent position of the nucleus at or outside the cell boundary in *vac14* PC3, σ-2. We visualize both sides of a PC axis to illustrate the features that are statistically linked. We interpret these results qualitatively as the sets of co-varying features that account for large proportions of statistical variation in the population. We found that while size scaling determines most of the morphological variability in the population, the relationship between organelle positioning, distance, shape, and orientation accounts for the rest. This is true in both normal and enlarged vacuole contexts, suggesting that organelle packing and confinement, arising from the distance or contact between organelles and occupancy of available space, is an important factor involved in determining shape and positioning, independently of or in parallel with size fluctuation.

### Biophysical interplay between vacuole mechanical properties and nucleus shape competition

Which physical properties of the vacuole and nucleus best explain the mechanical nature of steric packing interactions that may link the morphological features suggested by the integrating modeling and produce the real configurations seen in our dataset? An interesting observation from the integrated shape modeling is that, compared to normal cellular contexts, aberrant vacuole enlargement (in the sense of vacuoles being larger than expected from cell volume following the standard scaling relation) introduces more extreme effects on nuclear position and shape in addition to correlated vacuole-nucleus interfacial flattening. This indicates that abnormal degrees of vacuole-nucleus steric packing push nuclear morphology beyond normal ranges with only minimal effects on vacuole shape. In contrast, normal cells may have neutral geometry at the vacuole-nucleus interface, or in some cases feature a slight indentation on the vacuole. This asymmetric organelle relationship suggests “shape competition”, where the direction of deformation depends on cellular context and the relative physical or mechanical properties of the organelles in contact.

To examine the role of relative mechanical properties between the nucleus and vacuole and how this might modulate the effect of vacuole enlargement on the nucleus size, we systematically varied the mechanical properties of the vacuole in our biophysical model, by increasing the ratio of the vacuole to nuclear bending rigidity. We chose to increase the bending rigidity of the vacuole relative to the nucleus, a quantity we termed 𝛽, while keeping the surface tension constant given the experimental observation that vacuoles appeared more spherical as they enlarged. We varied the bending rigidity as 𝛽𝜅_#_, where 𝛽 = 1, 10, 50, 100. Increasing the vacuole bending rigidity relative to the nucleus led to larger vacuoles consistent with the combined action of osmotic pressure and relaxation of the bending energy driving outward expansion. Notably, simulations with more rigid vacuole membranes showed progressively flatter and smaller nuclei with increasing vacuole size (Fig. 6A).

**Figure 6.**
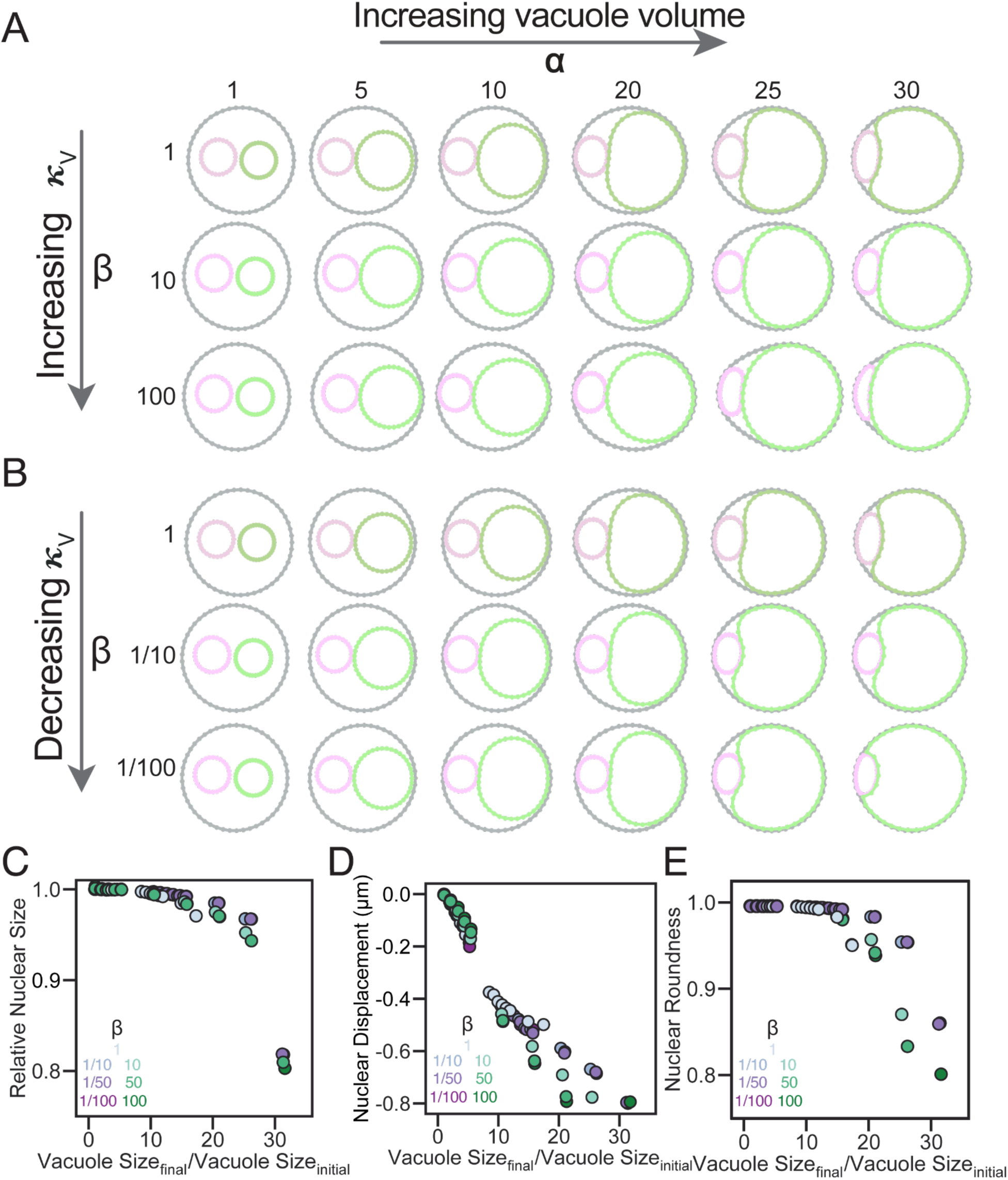
Comparing the impact of vacuole mechanical properties on nuclear morphology. Snapshots showing the final equilibrium morphologies of the nucleus, vacuole and cell after minimization using conjugate gradient varying the vacuole enlargement (left to right) and A) increasing vacuole bending rigidity (top to bottom). B) Decreasing vacuole bending rigidity C) Relative nuclear size, D) nuclear displacement, and E) nuclear roundness as a function of vacuole size and vacuole bending rigidity. Cases where vacuole bending rigidity is lower than nucleus are shown in magenta. Cases where vacuole bending rigidity is higher than the nucleus are shown in green.

Considering scenarios where the vacuole is softer, we repeated our simulations with𝛽 = 1, 1/10, 1/50, 1/100. Snapshots of the equilibrium morphologies show that at larger vacuole sizes, the vacuole indents and deforms around the nucleus (Fig. 6B). Quantification confirmed that softening the vacuole membrane also produced nuclear shrinkage and displacement, mirroring the effects of increased vacuole bending rigidity (Fig. 6C, D and E). However, for a comparable vacuole size, a softer vacuole membrane produced smaller reductions in nuclear roundness, area, and displacement than a stiffer membrane. Here, a softer vacuole membrane accommodates deformation preferentially within the vacuole itself, relieving the spatial constraint on the nucleus and preserving nuclear shape. Taken together, these observations indicate that the relative mechanical properties of the vacuole and nucleus govern the extent to which vacuole enlargement disrupts nuclear size scaling.

### Vacuole-nucleus proximity is a determinant of nuclear morphological dependence on vacuole size

One of the features highlighted by our integrated modeling results was that the spacing between the vacuole and nucleus varies with organelle shape, positioning, and spatial orientation (Fig. 5F, PC3 and PC4). Intuitively, packing effects require direct contact between objects. Could inter-organelle proximity be a more direct metric than vacuole volume ratio determining the impact of vacuole enlargement on nuclear morphology? Furthermore, could the degree of proximity distinguish between the roles of contact site tethering and steric packing in determining the vacuole-nucleus morphological relationship?

We categorized the entire dataset by vacuole-nucleus proximity status. If directly-adjacent vacuole and nucleus pixels could be detected, the organelles were considered to be in contact (“Contact” group). A minimum space of 1 or 2 pixels between the organelle boundaries was considered to be within the potential range of NVJ formation (“Small Gap” group, 30-60 nm). The separation of these categories is intended to separate steric packing, which should be characterized by the closest possible proximity between organelle membranes, from tether-based physical contact, which would be limited by tether length within the range of 30-60 nm (Eisenberg-Bord et al., 2016; Hariri et al., 2018), although these distances may not be precise in our data as they approach the limits of our spatial resolution. Thus, while both groups may include NVJ tethering, only the Contact group should reflect direct apposition of organelle membranes. A minimum space of larger than 2 pixels was designated as a lack of contact between organelles (“No Contact” group) (Fig. 7A, S4A).

**Figure 7.**
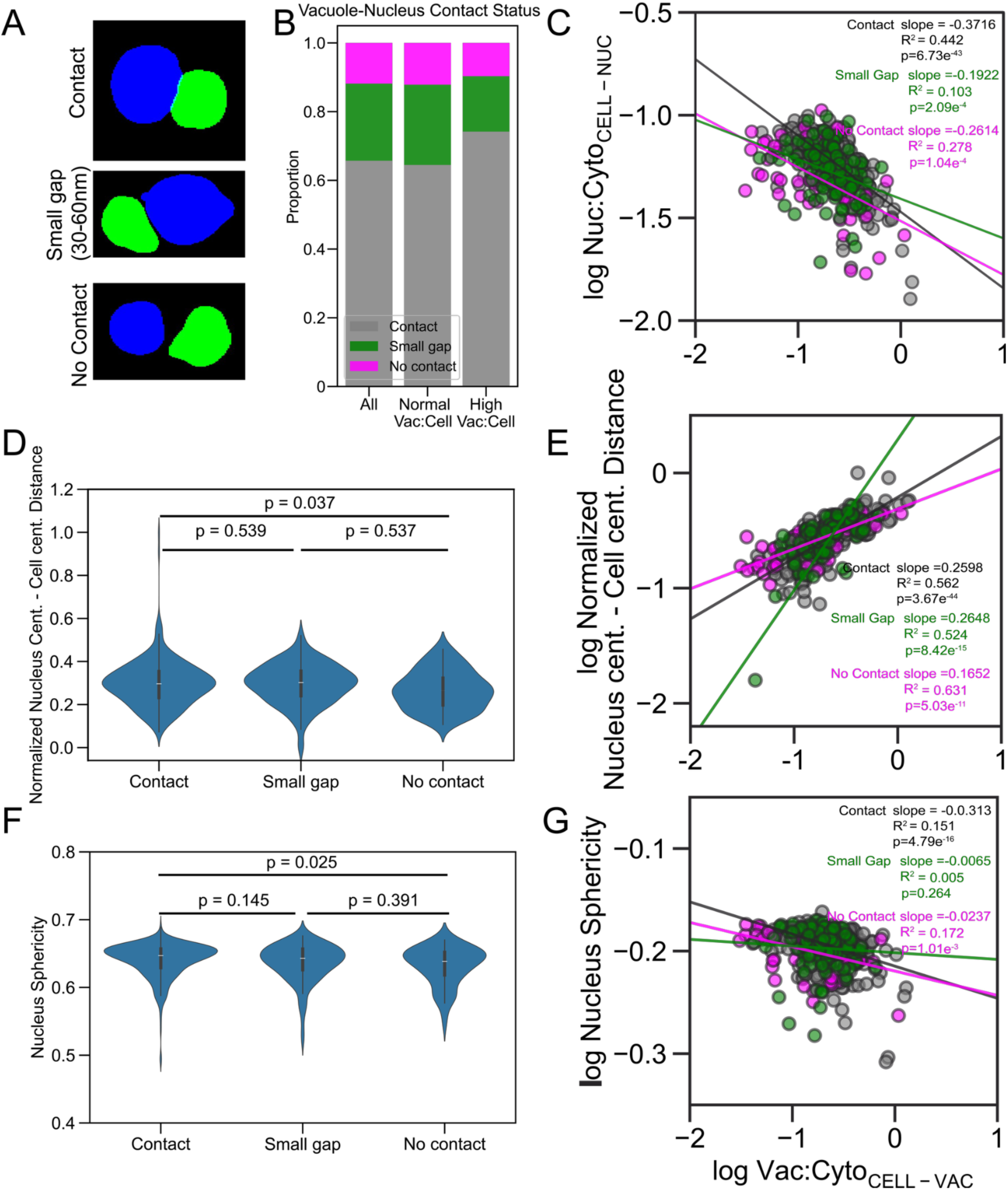
Vacuole size-driven effects on nucleus depend on vacuole-nucleus proximity. **A)** Voxel-based vacuole-nucleus contact status categorization. Vacuoles (blue) and nuclei (green) with direct pixel-pixel proximity were labeled “Contact”. Those separated by 1 or 2 pixels (30-60 nm) were labeled “Small Gap”. Those with a distance of at least 3 pixels were labeled “No Contact”. **B)** Proportion of cells by contact status for all 490 cells, normal V:C, and high V:C groups. **C-G)** Relationship between vacuole-nucleus contact status and vacuole size-dependent nuclear phenotypes: **C)** Nucleus:Cytoplasm volume scaling relationship with Vacuole:Cytoplasm. **D, E)** Nuclear positioning relative to the cell center. **F, G**) Nuclear sphericity.

As expected, most cells had vacuoles and nucleus in close proximity, with the majority in direct contact (322/490, 65.7%) and a large subset with a 30-60 nm gap (110/490, 22.4%). Interestingly, nearly 12% of cells had no vacuole-nucleus contact (58/490, 11.8%). We found a higher fraction of direct contact and lower fraction of “small gap” and “no contact” cells in high V:C cells compared to normal V:C cells (Fig. 7B, Fig. S4C). A similar difference was observed in *vac14* compared to VPH1-GFP (Fig. S4A-B), although there was no apparent difference in “no contact” proportion between these strains. Nuclear size scaling with V:Cyto is strongest in the Contact group (slope = -0.37, r^2^ = 0.442). Nuclear displacement is observed more frequently in the Contact group, with slightly higher average normalized nuclear distance from the cell center in Small Gap cells than No Contact (Fig. 7D), and a stronger scaling of distance with V:Cyto in the Contact group compared to the other groups (Fig. 7E). Nuclear sphericity is also related to contact status by central tendency and correlation with V:Cyto, with a significantly stronger effect in Contact cells (Fig. 7F-G). Unexpectedly, the scaling relation for nuclear volume ratio and sphericity with V:Cyto is slightly stronger in the No Contact group (slope = -0.26, r^2^ = 0.278) than in the Small Gap group (slope = -0.192, r^2^ = 0.103) (Fig. 7C and 7G).Taken together, these results show that all nuclear phenotypes are stronger when the vacuole and nucleus are in direct contact.

To further explore the interplay between contact site tethering, sterics, and organelle driven packing on nuclear morphology, we introduce tethering, modeled as springs between the nucleus and the vacuole into our computational framework. Here, when a node on the nucleus is within a cutoff distance of an edge of the vacuole, a tether is formed between the nucleus node and the closest point to the nucleus node on the vacuole. The cutoff distance we chose is 30 nm, in line with the distance of an NVJ tether. Due to the lack of available literature on the rates governing the dynamics of tether formation, we chose to model tethering as a binary switch and without accounting for tether dissociation. Once formed, tethers remain attached for the remainder of the simulation.

We varied the size of the vacuole, the mechanical properties of the vacuole and the presence or absence of tethering (+/- tethering) and examined the nuclear morphology. Our simulations reveal that the relative bending rigidity of the vacuole and nucleus governs how tethering reshapes the nuclear envelope. When the two organelles have equivalent bending rigidities, tethering at the contact site produces a localized dimple in the vacuole as the nucleus presses against and deforms the vacuole in contrast to the case without tethering (Fig. S5A). Increasing the vacuole bending rigidity reverses this asymmetry: the nucleus instead deforms around the stiffer vacuole; tethering drives a transition from a spherical to an ellipsoidal nuclear shape that becomes flatter as the vacuole enlarges (Fig. S5A). Decreasing the vacuole’s bending rigidity, by contrast, produces little change in nuclear morphology across the range of vacuole sizes examined. Quantitative analysis confirms these trends, showing that an increasing ratio of vacuole to nuclear bending rigidity yields nuclei that are both flatter and more displaced from their original position (Fig. S5C, D). Nuclear size, however, plateaus as the vacuole becomes stiffer, reflecting the spreading of the nucleus along the tethered surface. This suggests that the mechanical properties and volume constraints of the nucleus limit how far the nucleus can extend (Fig. S5B). For softer vacuoles, tethering exerts no discernible effect on nuclear shape, with the bending rigidity curves collapsing onto a single curve (Fig. S5 E, F, G). We note that these simulations model tethering as a binary on/off interaction and therefore capture the extreme limits of its mechanical influence; even so, the results suggest that nuclear geometry may serve as a useful indicator for distinguishing tethered from non-tethered states.

### High-resolution whole cell geometric and morphological analysis illustrates distinct features of organelle packing

The differences in results between Contact and Small Gap groups suggest either that packing effects in the vacuole-nucleus interaction are distinct from that of contact site tethering without sterics, or that an NVJ is not necessarily present even when the organelle membranes are within 60 nm of each other. What is the relationship between tethering and packing of organelle membranes? Can organelle membranes be closely packed due to steric constraints without necessarily forming a contact site, or is proximity always accompanied by tethering? If so, can packing induce contact site adhesion by bringing membranes in close enough proximity to form stable tether complexes, or is an initial contact site first required to enable close apposition of membranes which can in turn lead to expansion of the contact site upon stronger membrane packing? Can organelle packing as a result of either vacuole expansion or boundary constraint from organelle orientation relative to cell wall diameter, as shown in Fig. 5, drive similar morphological effects on organelles, or are these two conditions typically linked?

We selected individual instances of cells representing distinct geometric conditions for complete morphological characterization of organelle shape, positioning, and interface geometry (Fig. 8) The first example is a normal V:C cell with typical nuclear morphology and direct vacuole-nucleus contact (Fig. 8A-H). The organelles are not distributed along the cell’s long axis. The minimum distance between organelle membranes and the cell wall along the vacuole-nucleus (V-N) axis is similar or larger than the overall minimum distance along any axis, suggesting that there is sufficient free cytoplasmic space available for organelles to position normally without vacuole-nucleus packing effects enhanced by boundary constraint (Fig. 8A, C). Vacuole and nucleus shapes have similar values by voxel-based sphericity, reflecting typical shapes in the near-spherical range (Fig. 8B, D). The mesh curvature-based metric Willmore energy density (WED) is more sensitive, revealing a larger deviation from sphere shape for the vacuole (Fig. 8E), attributable to a large globular section emerging from the main body of the organelle (Fig. 8A). Vacuole-interface detection to delineate the regions of the vacuole and nucleus surfaces in direct contact and at distances consistent with putative contact site tethering (Fig. 8F, N, V). The majority of the total interface region is in direct contact, such that the mesh surfaces have no discernable distance between them (13.2% of the total nuclear surface area and 4.9% of the vacuole surface area, while a narrow surrounding region is within the 0-60 nm that could harbor a tether-based contact site (19.3% of the nuclear and 7.1% of the vacuole total surface area). (Fig. 8G, H).

**Figure 8.**
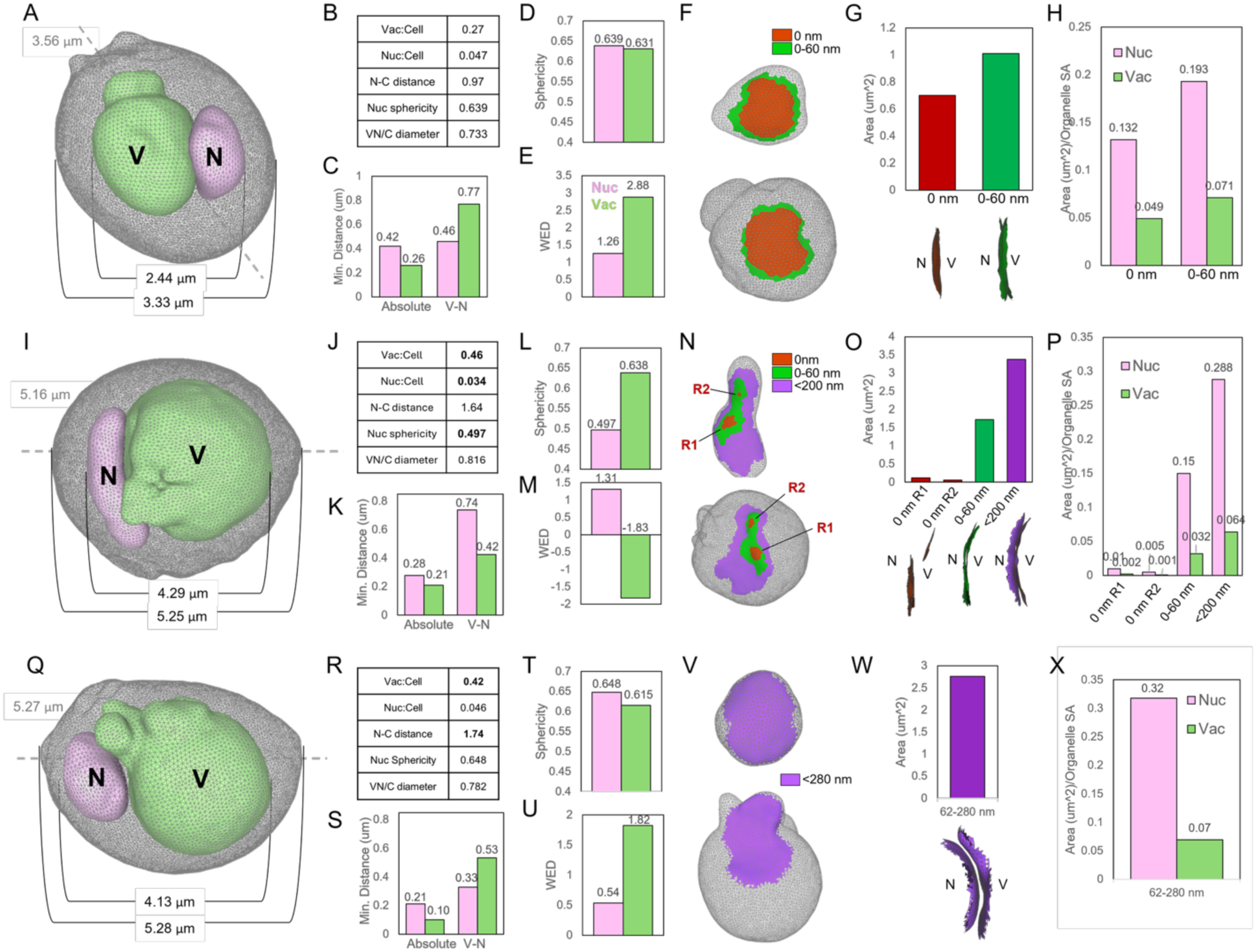
Whole-cell mesh representations illustrate distinct geometric states of vacuole enlargement and organelle interaction. **A-H)** Normal V:C cell with normal nuclear morphology and direct vacuole-nucleus contact. **I-P)** High V:C cell with low N:C, flattened nucleus, and direct vacuole-nucleus (V-N) contact. **Q-X)** High V:C cell with displaced nucleus and no V-N contact. **A, I, Q)** Whole-cell view of cell (gray), vacuole (green), and nucleus (pink) reconstructed as triangulated meshes. Measurements indicate joint V-N diameter (inner bracket), cell diameter along V-N axis (outer bracket), and cell diameter along long axis (gray dashed line). Ratio of V-N diameter to parallel cellular diameter (VN:C) is indicated. **B, J, R)** Basic statistics including the vacuole:cell (“Vac:Cell”) and nucleus:cell (“Nuc:Cell”) volume ratio, normalized nucleus-cell centroid distance (“N-C distance”), and nuclear sphericity (“Nuc sphericity”). **C, K, S)** Minimum distance of organelle surfaces to cell boundary, measured as absolute minimum distance at any point (“Absolute”) or along V-N axis (“V-N”). Organelle shape quantification by **D, L, T**) voxel-based sphericity and **E, M, U**) Willmore energy normalized by surface area (Willmore energy density (WED)). **F-H, N-P, V-X)** Vacuole-nucleus interface analysis. **F, N, V)** Interface mapping on nucleus (top) and vacuole (bottom) surfaces at multiple distance thresholds (0 nm (red), 0-60 nm (green), and purple region represents full manually delineated interface area, with max distance indicated. **G, O, W)** Surface area as an average of the vacuole and nucleus component of each interface. Side-view of each mesh-rendered interface region is displayed below. **H, P**, **X)**. Interface area fraction of total organelle surface area for each interface region.

The second example represents a typical high V:C cell with a strongly flattened nucleus and organelles in contact (Fig. 8I, J). Here, the vacuole-nucleus axis is aligned parallel to the cell’s long axis, occupying a larger proportion of the cell diameter along this axis compared to the first cell (81.6% vs 73.3%). Although this cell clearly exhibits packing effects, there is a similar degree of space between the organelles and the cell boundary at the vacuole-nucleus axis, suggesting a limit to proximity between organelles and the cell wall, possibly due to the presence of other structures we cannot see (Fig. 8K). Although the nucleus is flatter than the vacuole, the vacuole exhibits irregular shape features including dimples possibly indicating lipid droplet positioning (Chen, et. al. 2025), and protrusions extending around the nucleus (Fig. 8A, L, M). Interface detection reveals only two small regions in direct contact and a substantial region of 0-60 nm MCS-compatible proximity. However, tracing the full surface area in which the two organelles exhibit apposition with corresponding curvature, such that the vacuole membrane is convex where the nucleus membrane is concave from the side view shows that physical interaction between the organelles does not exclusively occur where there is direct membrane-membrane contact (Fig. 8N, O, P).

The final cell represents a rare case in which the vacuole is enlarged (Vac:Cell 0.42), but the nucleus and vacuole are not in contact with each other (Fig. 8Q). This example illustrates the role of tethering or direct contact in organelle total and/or interface geometry when organelles are packed. In this case, the Vac:Cell ratio is similar to that in the previous case, but no nuclear size or shape phenotype is observed, although the nucleus distance from the cell center indicates displacement (Fig. 8R). Similarly to the previous case, organelles are positioned parallel to the cell long axis while maintaining a small distance from the cell wall, and the vacuole has a large sphere-like section as well as an irregularly shaped section extending around the nucleus (Fig. 8Q, T, U). The entire interface region is outside of the range of NVJ proximity (62-280 nm) with a large area covering approximately one-third of the nuclear surface area (Fig. 8V, W, X). In contrast with the previous example, the curvature correspondence between membranes is convex on the nuclear side and concave on the vacuolar side. These observations suggest that, while direct contact is not necessary for packing-driven local deformations, the directionality of deformation, namely nuclear flattening and indentation, appears to require direct inter-organelle contact.

## Discussion

In this study, we combined detailed analysis of the vacuole-nucleus morphological relationship through high-resolution, high-throughput 3D whole-cell imaging coupled with linear modeling and biophysical simulations to reveal how steric packing effects of vacuole enlargement impact the nucleus. Our multifaceted quantitative approach represents a framework to link static statistical morphological information with physics, isolating the role of mechanical interactions on morphology from dynamics and molecular regulatory mechanisms. We were thus able to quantitatively characterize the precise effects of vacuole enlargement on nuclear features, as well as identify covariations between features and their explainability by simple physical mechanisms.

The main organelle features we studied, namely size control, subcellular positioning, and shape, have each been extensively characterized as important biological properties with distinct biological functions, dynamics, complex mechanisms of regulation, and roles in disease. Our work examines an understudied unifying aspect - the variability within as well as outside of normal ranges in each parameter as a secondary consequence of dysregulation in another organelle, the vacuole in this case, rather than due to intrinsic behavior of or direct regulation on the nucleus by active mechanisms. Nuclear size scaling, also known as nucleocytoplasmic or karyoplasmic ratio, is considered to be a generally constrained parameter across eukaryotes (Gillooly et al., 2015). However, we found that not only can the nucleocytoplasmic ratio become altered when vacuoles are enlarged, but this loss of normal nucleocytoplasmic scaling is well tolerated by cells, and is in fact seen at relatively low degrees of vacuole enlargement (Figures 1 and 4).

These results support two main conclusions. First, vacuole size impacts distinct features of the nucleus in a size-dependent manner: nuclear size scaling is affected first, followed by displacement, and finally nuclear flattening only occurs at high vacuole sizes. Second, when the vacuole is enlarged above normal range, the effect on nuclear size ratio and position is reduced, possibly indicating a limit to the tolerable effect on these parameters, while the effect on nuclear shape is more severe. These observations suggest that there is a range of adaptive nuclear size and positioning in healthy cellular regimes that contributes to normal morphological variation with no associated dysfunction. In contrast, nuclear shape is generally preserved until vacuole size increases to an abnormal degree, suggesting that nuclear deformation is a more dysfunctional effect.

The observation that the nucleus is displaced from the cellular center toward the periphery in relation to vacuole size even within normal vacuole size ranges suggests that nucleus position is a variable parameter that responds to packing interactions in normal regimes. Thus, packing interactions are present to some degree in normal cellular conditions, not only in dysfunctional states that cause the abnormal enlargement of organelles or crowding by other means.

Nuclear shape, on the other hand, is only significantly affected at higher degrees of vacuole enlargement. The relative thresholds of vacuole enlargement or spatial constraint required to perturb each nuclear property may reflect the cell’s relative tolerance to variation or disruption in that property. On one hand, the nucleus is mechanosensitive and its shape is closely associated with chromatin organization, and nuclear deformation is an important marker of aging (Belhadj et al., 2023) and cancer (Dey, 2009). On the other hand, nuclear deformability can play important roles in processes such as closed mitosis in yeast, when the nucleus undergoes constriction and drastic shape transitions in order to pass through the bud neck. Matos-Perdomo and colleagues recently showed that yeast nuclei form irregular toroidal shapes scaffolded by the vacuole to accommodate rDNA loop formation in mitosis (Matos-Perdomo et al., 2022; Santana-Sosa et al., 2023). This represents a functional cellular context for a morphological vacuole-nucleus interaction independent of NVJ tethering. Interestingly, it was proposed that the nuclear shape remodeling could be a mechanism to regulate nucleocytoplasmic ratio as the rDNA expands.

The 3D spatial information offered by whole-cell reconstructions is ideal for revealing packing interactions between organelles, although these interactions are not commonly directly quantified or deliberately studied. Several whole-cell imaging studies have qualitatively observed inter-organelle morphological relationships indicating packing phenomena (Ferreira LR et al., 2008; Loconte et al., 2021; Matharu SS et al., 2023). These observations in distinct biological contexts, in combination with our study, raise the question of whether organelle packing is a universal or context-dependent phenomenon in cells. Standardized morphological hallmarks and quantitative measures of organelle packing are needed to evaluate the degree, cross-organelle asymmetries, and underlying causes of organelle packing, as opposed to general cytoplasmic crowding or density (Loconte et al., 2022). A caveat to this approach is that most image datasets will not be able to provide volumetric information for all contents of the cytoplasmic space, so any given designation of “available cytoplasm” must be considered to contain an unknown composition of other cellular structures in addition to truly “available” fluid space. Furthermore, volume ratios alone do not reflect true features of organelle packing such as the shape relation between packed organelles resulting from direct physical contact and mechanical forces. Our spatially resolved mechanical model demonstrates that vacuole enlargement alone, through excluded-volume confinement and the relative mechanical stiffness of the vacuole membrane, is sufficient to disrupt nuclear size scaling, positioning, and shape in a size-dependent manner, providing a minimal physical mechanism for inter-organelle morphological interaction (Figs. 4 and 6). Future progress in organelle packing studies will benefit from the development of a crowding metric that holistically reflects the degree of morphological constraint on organelles. This effort should consider the utility of organelle packing as a cell-level metric (i.e., the degree to which the cell cytoplasm is filled with organelles), in contrast with an organelle-level metric (i.e., the degree to which each organelle is constrained by packing from other organelles). Variation between organelles in the same cell in this latter metric could reveal hierarchical physical interactions between organelles, yielding insights into the overall physical interplay and homeostasis of the organelle interactome.

The choice of budding yeast as a model system presents several advantages. High numbers of *S cerevisiae* cells can be imaged and tomographically reconstructed by SXT thanks to their small size relative to the SXT capillary, and the strong precedent of SXT applied to budding yeast including well-established organelle LAC values used as a baseline for segmentation (Chen et al., 2025; Larabell & Le Gros, 2004; Uchida et al., 2011). The simple, rigid shape of cell wall-bound yeast cells and the relatively simple globular shapes of nuclei and vacuoles, along with straightforward synchronization or visual screening for unbudded cells in S. cerevisiae cells allows us to control multiple sources of morphological and biological variation in the cell population, facilitating geometric modeling of the vacuole-nucleus relationship with interpretable covariants. Together, these features enable the translation of well-understood genetic conditions to a high-precision phenotypic modeling, which allows us to understand a specific geometric cell state (enlarged vacuole) in terms of its cell-wide effects. SXT-derived surface meshes of whole yeast cells now enable precise 3D curvature analysis of entire organelles at high throughput, and integrating such structural data with the biophysical properties provided by SXT offers a promising avenue for holistic whole-cell modeling. A natural extension of the present modeling framework is to incorporate experimentally derived organelle geometries to replace idealized initial conditions (Lee, Laughlin, Beaumelle, et al., 2020; Lee, Laughlin, Moody, et al., 2020; Zhu et al., 2022), enabling single-cell mechanical predictions to be validated against population-level morphometric distributions.

On the other hand, the generalizability of our approach is limited in other systems that don’t share this set of characteristics. The intrinsic degree of cell-to-cell variability, even in cell populations synchronized for cell cycle stage, cell shape, and environment, may be higher in other cell types or even other strains of budding yeast. The dataset size needed to benchmark an integrated model should represent the baseline variability in the population, which will present technical challenges to producing datasets of sufficient spatial resolution large enough to represent more variable cell types. Our dataset is relatively uniform and yet requires a large sample size for the technique and resolution to yield an informative integrated model. Feasibility can be increased with strategic choices of combinations of parameters including cell state and environmental controls, cell subset selection, target organelle identity and number, imaging resolution, cell alignment axis, number of spherical harmonics coefficients used to reconstruct shapes, or other shape modeling schemes such as that may be better suited, as spherical harmonics expansion only works on sphere-like, single copy organelles (Johnson et al., 2015; Keren et al., 2008; Li et al., 2016; Pincus & Theriot, 2007). The present model’s treatment of the system as a deterministic mechanical equilibrium omits active fluctuations and cytoskeletal force contributions that are known to modulate nuclear deformation in other contexts (Doolin et al., 2019), and future iterations incorporating thermal, active, and biochemical fluctuations, such as stochastic cytoskeletal remodeling, would enable direct comparison of simulated morphological variance with the cell-to-cell variability observed experimentally (Monfared et al., 2025), thereby transforming the present deterministic model into a probabilistic framework capable of predicting population-level morphological distributions.

## Materials & Methods

### Yeast image data acquisition and analysis

#### Yeast soft X-ray tomography imaging

The generation of the yeast soft X-ray tomography data analyzed here is described in detail in Chen, Mirvis, Ekman *et al* 2025 (Chen et al., 2025). Briefly, BY4741A (wild-type haploid, mating type a), BY4741A *vph1-GFP::his3*), VPH1-GFP (*vph1-GFP::his3* in BY4741A background), and *vac14* (BY4741A *vph1-GFP::his3 Δvac14*) were grown to early log cultures of OD600 0.2-0.4. Cells resuspended at a density of 10^6^ were prepared for soft X-ray tomography imaging. Resulting tomographic reconstructions were segmented into single cell volumes and segmented using a custom autosegmentation 3D U-Net model trained on a subset of manually segmented images. 3D multichannel segmentations were manually quality-checked and filtered to include only complete volumes of unbudded, early G1 cells with no signs of cellular stress or death.

#### Quantitative morphological analysis

Voxel-based morphometrics measurements (volume, surface area, centroid, Feret diameter) were produced using the 3D Suite package for FIJI (Ollion et al., 2013; Schindelin et al., 2012; Schmid et al., 2010). Statistical significance was tested for comparison of distributions by Mann-Whitney U-test for nonparametric distributions (Fig. S1, Table S1) using the Scikit stats Python package (*Numpy and Scipy Documentation — Numpy and Scipy Documentation*, n.d.).

Mesh creation and processing from binary stacks was performed in BlendGAMer (Lee, Laughlin, Beaumelle, et al., 2020; Lee, Laughlin, Moody, et al., 2020) and MeshLab (Cignoni et al., 2008) as previously described (Chen et al., 2025). Multichannel binary segmentation z-stacks were channel separated, and each channel was converted to a binary triangulated surface mesh and exported as an STL image using the FIJI 3D Volume Viewer plugin (Schmid et al., 2010). STL files were imported to Blender version 3.6 and scaled to bounding box dimensions measured in FIJI using the 3D Object Counter tool (Bolte & Cordelières, 2006). The BlendGAMer plugin, version 2.0.8 (Lee et al., 2024; Lee, Laughlin, Beaumelle, et al., 2020) was used for mesh smoothing using the following steps: smoothing (15 iterations), Coarse Dense decimation (threshold 1.5, 10 iterations), smoothing (15 iterations), Coarse Dense decimation (threshold 1.0, 10 iterations), smoothing (10 iterations), smooth normals.

Distance measurements including vacuole-nucleus joint diameters and cell diameters were taken manually using the Measurement tool. Five separate measurements were taken from different cell rotations and averaged to report a single value. Mean and Gaussian curvatures were calculated using the Discrete Curvature function. Average, standard deviation, and variance of curvatures were recorded and used to calculate Willmore energy (WE) density as previously described using the formula:

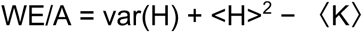

Where WE is Willmore energy, A is surface area, H is mean curvature, K is Gaussian curvature, and brackets denote average.

Interface detection was performed based on signed distance maps generated using the Distance from Reference Mesh function and selecting vertices by the distance thresholds described (Fig. 8, Conditional Vertex Selection function). Selected regions were converted to faces by manual tracing and moved to separate layers. Geometric and curvature measures were recorded for each interface region. Interface areas are reported as the average of corresponding vacuole and nucleus interface regions.

#### Integrated shape modeling

Integrated modeling code was installed from Allen Cell’s Github repository (https://github.com/AllenCell/cvapipe_analysis), and parameters were modified slightly to add spherical harmonic parameterization for the vacuole while omitting fluorescence mapping steps. Single cell images were loaded into the pipeline and smoothed using a Gaussian kernel σ=2. The analysis package was used to extract morphological features, including volumes and spherical harmonic coefficients using the Allen Cell spherical harmonics parameterization package (L_max_ = 16) (https://github.com/AllenCell/aics-shparam). Mitotic cells and feature-based outliers are removed from the dataset, and the remaining dataset is used to run principal component analysis. The resulting shape modes are then visually produced as 2D animated GIFs and VTK mesh files representing the average shape for each structure.

### Simulation Model and Assumptions

In our model, we treated the organelles as enclosed 2-D polygons with resistance to bending and stretching. The energy of the membrane is given by discrete 2-D Helfrich Hamiltonian,

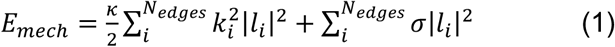

Where 𝜅 is the bending rigidity of the organelle membranes and 𝛔 is the surface tension, *k*ᵢ is the local curvature at vertex *i*, and *l*ᵢ is the vector of the edge connecting vertices *i* and *i*+1.

There are several established ways to define curvature in discrete settings. For this model, we choose a definition of curvature based on the turning angle at each vertex.

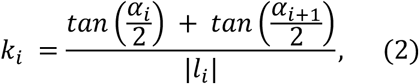

where the turning angle *α*ᵢ is given by the sum of the internal angles,

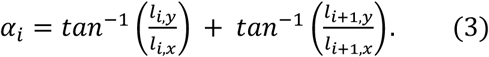

#### Steric repulsion between organelles

To account for excluded volume interaction between the organelle and cell wall compartments, we implemented a repulsive potential between the edges of the membrane polygons. We discretize each polygon edge into N evenly spaced points. The steric repulsive potential between a compartment i and j is given by

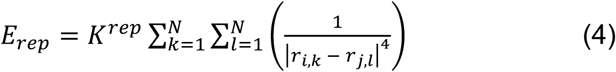

Where 𝐾*^rep^* is the repulsive strength, and r_i,k_ and r_j,l_ are the points on compartments i and j, respectively. The magnitude of repulsive energy interaction between edges of different polygons decreases rapidly with larger distances between the edges.

#### Organelle size control

To modulate the size of the organelles, we build on established theory showing that the osmotic conditions between the nucleus and the cell determines the steady state size of the organelle (Lemière et al., 2022b) where the osmotic pressures, 𝛥𝑃 are balanced by the mechanical forces in each membrane. In our model, this shows up as a pressure-volume work exerted on the organelle membrane taking the form

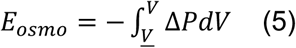

where <u>𝑉</u> represents the preferred volume of the organelle. To grow and shrink the organelles, we assumed the Van’t Hoff law to satisfy osmotic pressure given by

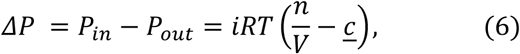

where *i* is the van’t Hoff index, R is the ideal gas constant, T is the temperature, n is the number of solute particles enclosed in the organelle, V is the volume of the organelle compartment and 𝑐is the ambient molar concentration.

For the cell, <u>𝑐</u> is the external concentration outside the cell, 𝑐*^out^*:. For the nucleus and the vacuole, <u>𝑐</u> is the cytoplasmic concentration, 𝑐*^cytoplasm^* Putting 𝛥𝑃 into equation (5) and integrating gives

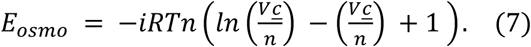

We can rename 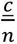 as <u>𝑉</u> since it denotes the preferred volume of the organelle and cell cytoplasm given the internal and ambient osmotic conditions. Here, the pressure-volume energy is determined by the ratio of the organelle volume and the preferred volume of the organelle. So the closer the organelle volume is to the preferred volume, the smaller the osmotic energy contribution. We group 𝑖𝑅𝑇𝑛 into one variable, *K_v_* which we term the osmotic strength constant. Since our framework is two-dimensional, we require a way to transform between area and volume required for the osmotic energy term. To achieve this, we assumed a linear scaling between the organelle areas and their volumes. Assuming a spherical organelle, the volume is given by 4/3πr^3^ and the cross-sectional area is given by 𝜋r^2^.

Assuming a change in the radius by dr, the volume-to-area ratio is given by

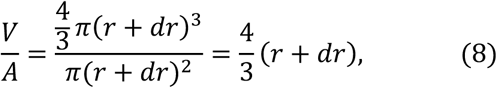

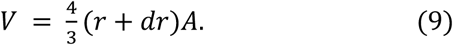

We approximate 𝑟 + 𝑑𝑟 as the average distance from each node in the polygon to the polygon centroid. This serves as the scaling factor between the organelle area in our 2-D framework and the volume in the osmotic energy.

#### Tethering between Organelles

We treated tethers as springs and the energy of the tether is given by

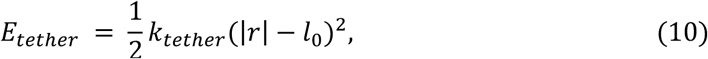

Where 𝑘*_tether_* is the tether spring constant, r is a displacement vector of the tether, and 𝑙_0_ is the equilibrium tether length. To form a tether, we find the closest edge on the vacuole polygon to each node on the nuclear polygon. If the distance between the nucleus and the vacuole is smaller than a cutoff distance, we place a tether between the nucleus node and the vacuole edge. Once a nucleus node and vacuole edge are tethered, those nodes and edges are removed from the available pool of points that can be tethered.

### Energy minimization and force calculation

To examine steady-state configurations of nucleus, vacuole and cell, we propagate the system to mechanical equilibrium by minimizing the energy of the system using the conjugate gradient method. The energy of each organelle *i* and the cell is given by

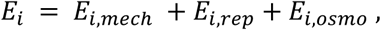

and the forces on each compartment *j* is given by

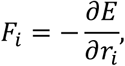

where 𝐹*_i_* is the force on the organelle *i* from the membrane mechanics, osmotic pressure, and steric interactions. The forces are calculated as the negative gradient of the energy with respect to the organelle coordinates r_i_. The system is simulated until the following stopping conditions: 1) magnitude of the gradient is less than 0.01 or 2) the value of the gradient does not change for 500 iterations.

#### Simulation setup

To investigate the effect of vacuole enlargement on nuclear morphology, we set the initial organelle and cell sizes using experimentally reported values of organelle sizes. We systematically varied the target size of the vacuole and the vacuole bending rigidity which yielded a range of enlarged vacuole configurations. We assume that the nuclear volume and cell cytoplasm volume, which is the cell volume less the nucleus and vacuole volumes, do not undergo any swelling or shrinkage from osmotic pressure, i.e., the final target volume (<u>𝑉</u>) is equal to the initial volume. We refer the reader to Table 2 for a detailed list of simulation parameters. We simulate the system using the conjugate gradient scheme until the energy difference between iterations is less than 1e-5. Further analysis is carried out on the resultant equilibrium organelle and cell morphologies.

## Acknowledgements

The soft X-ray tomography dataset used for this analysis, described in reference (Chen et al., 2025), was developed in collaboration with Carolyn Larabell as well as members of her group, especially Jianhua Chen and Axel Ekman. We thank them for extensive discussions about both SXT methods and mesoscale structure of cells. We also thank current and past members of the Marshall lab and Padmini Rangamani for many interesting conversations about this work and about the general question of organelle packing, Mark Chan for many discussions about yeast vacuole size, Jennifer Fung for the gift of yeast strains, and Matheus Viana and Susanne Rafelski at the Allen Institute for Cell Science for their invaluable help guiding us in using the integrated modeling software. Simulations were run on hardware hosted by the Triton Shared Computing Cluster (San Diego Supercomputer Center, 2022). We are also grateful to the UCSD Physics Computing Facility for computational resources. This work was supported by NIH grant R35GM130327 and by the Center for Cellular Construction, supported by NSF grant DBI-1548297.

---

**Figure S1.**
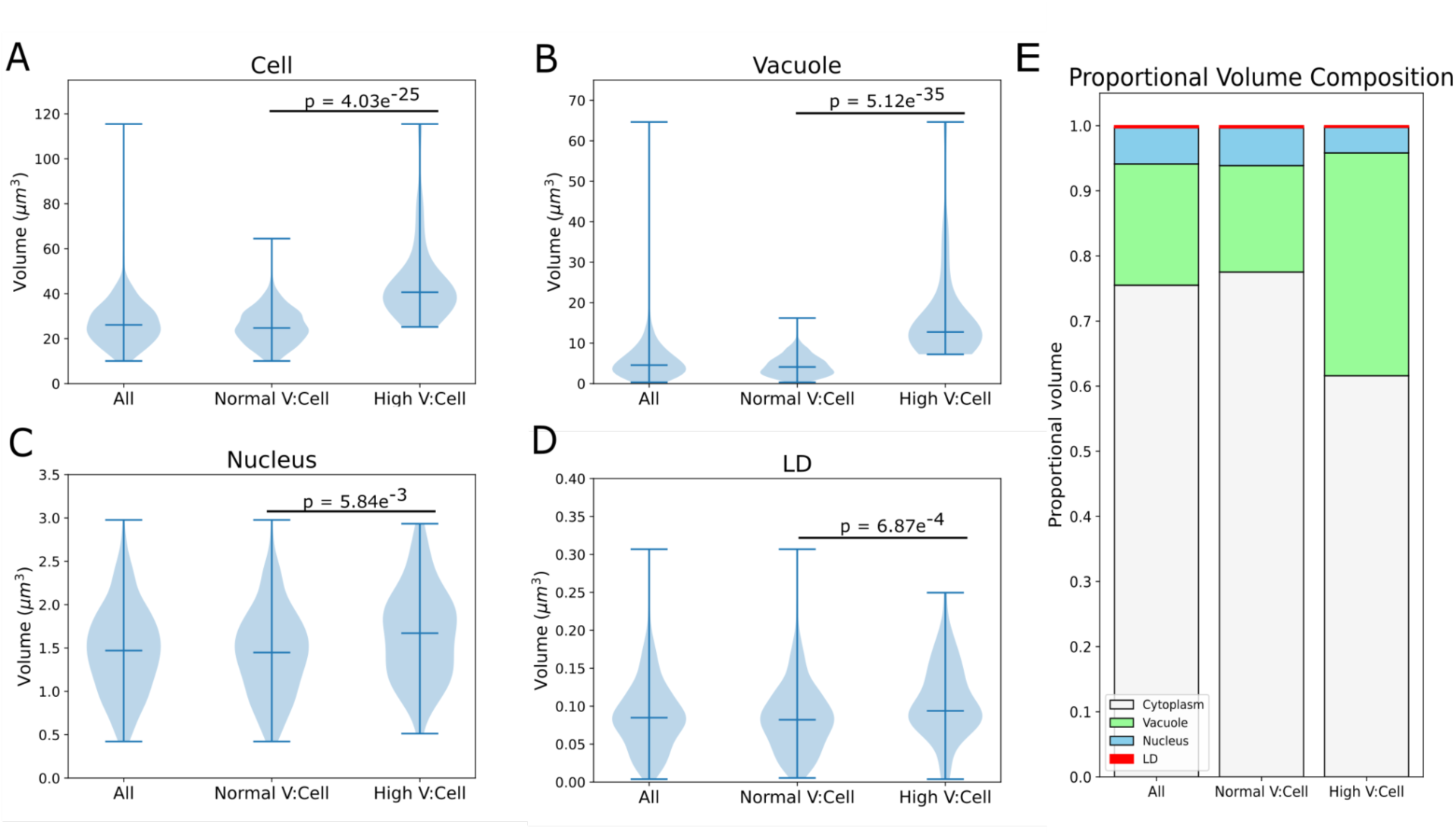
Cell and organelle volume distributions for normal and large-vacuole cell groups. Measurements taken from voxel-based 3D binary segmentations of reconstructed SXT volumes. A-D) Cell, vacuole, nucleus, and lipid droplet (LD) volume distributions for the pooled dataset, as well as “normal” and “high” V:C subsets divided at a threshold of 0.27 V:C based on voxel measurements of volume from SXT whole-cell reconstructions. Statistical significance was tested by Mann-Whitney U-test for nonparametric distributions (see Table S1). E) Normalized volume composition for each group. “Cytoplasm” is defined as the remaining volume after subtracting known compartment volumes (vacuole, nucleus, LD) from cell volume.

**Figure S2.**
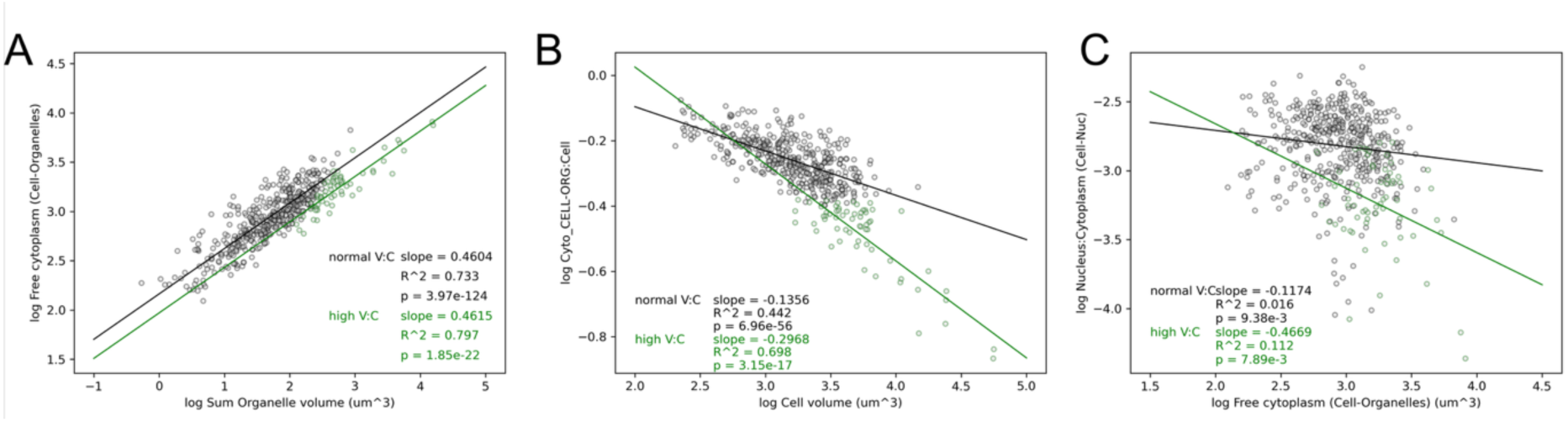
Log-log scatterplots describing scaling relations of free cytoplasmic space, defined as the cellular volume not occupied by known organelles (“free cytoplasm”, “Cell-Organelles”). **A)** Free cytoplasm volume scaling with total known organelle volume (“Sum Organelle volume (um^3^)). **B)** Proportional free cytoplasm volume (“Cyto_CELL-ORG_:Cell”) scaling with cell volume. **C)** Nucleocytoplasmic ratio (“Nucleus:Cytoplasm (Cell-Nuc)”) scaling with free cytoplasm (“Free cytoplasm (Cell-organelles) (um^3^)).

**Figure S3.**
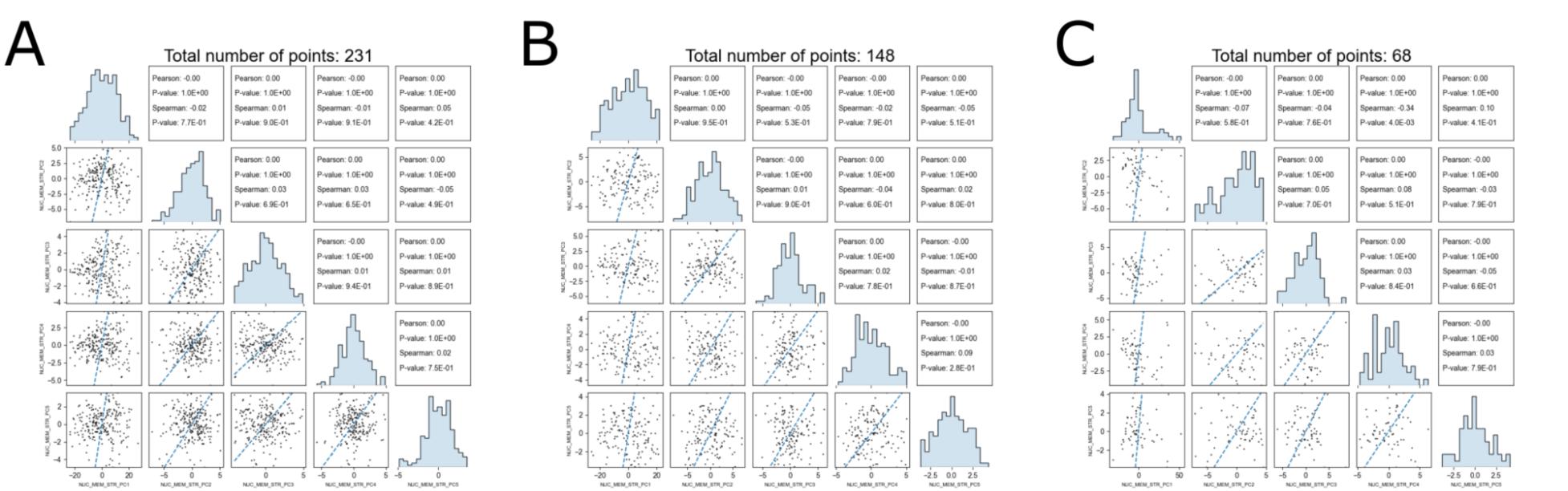
Pairwise correlations of principal components from integrated shape modeling for **A)** WT, **B)** VPH1-GFP, and **C)** *vac14* strains. Central histograms show smooth, generally unimodal population of the shape space for each PC. Each PC combination shows no significant correlation (>1E-2) by Pearson or Spearman tests, indicating separation of distinct principal components.

**Figure S4.**
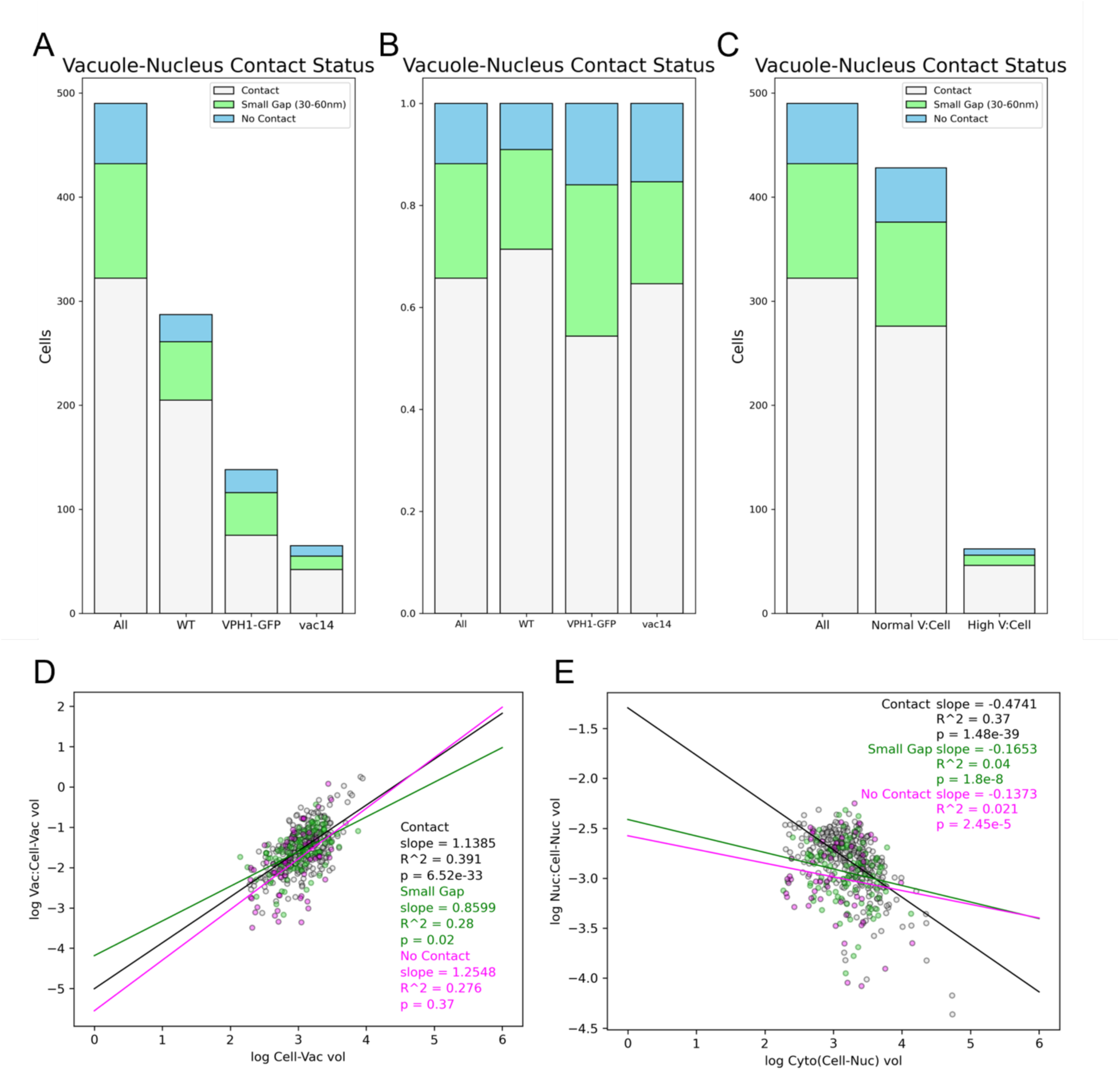
Vacuole-nucleus proximity. A) Raw number and B) proportion of cells by contact status for all 490 cells, and separated by genetic strain. C) Raw number cells by contact status for all 490 cells, normal V:C, and high V:C groups. D) Vacuole:Cytoplasm and E) Nucleus:Cytoplasm size scaling with cytoplasmic volume (excluding vacuole or nucleus, respectively) based on contact group.

**Figure S5.**
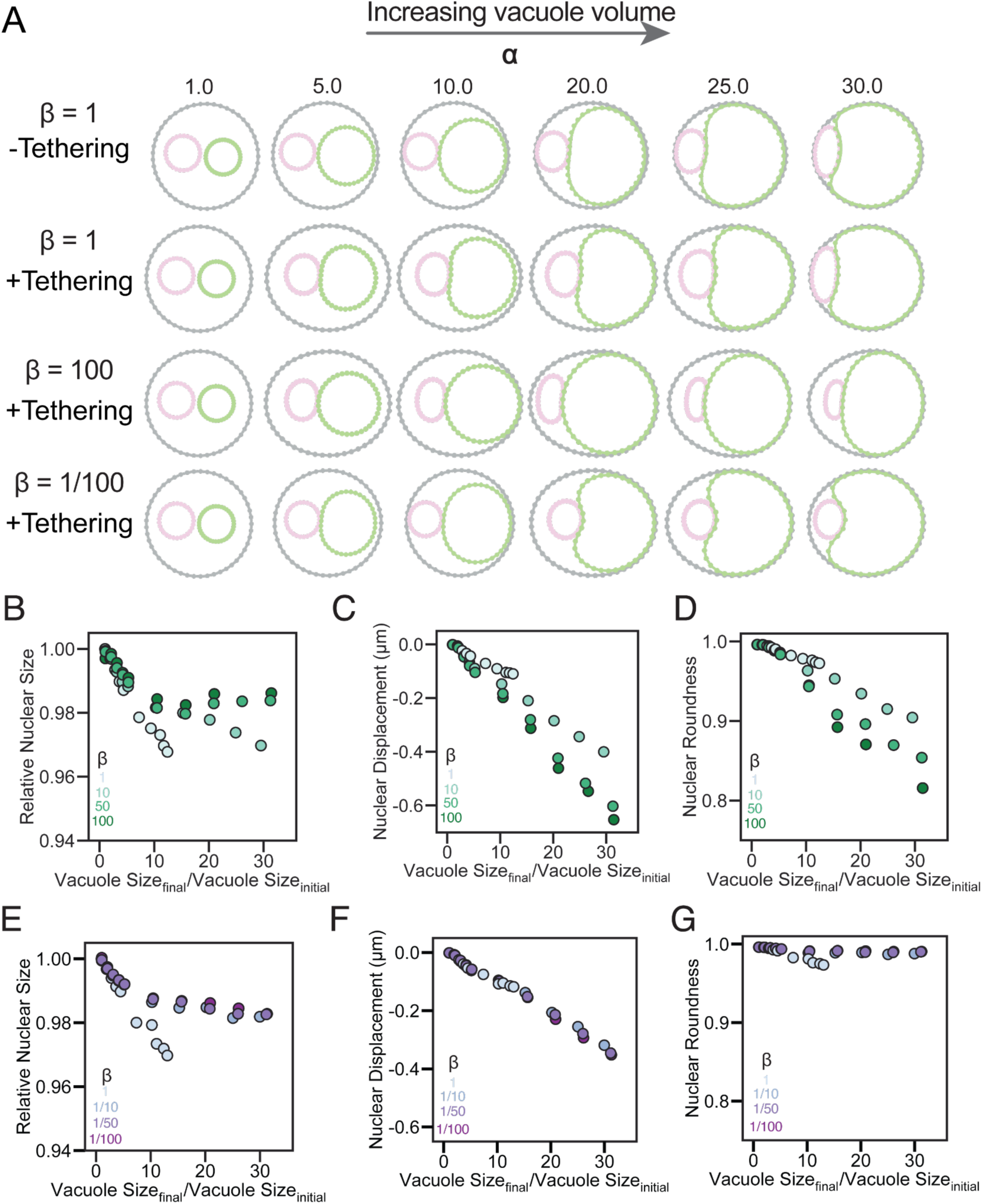
Impact of tethering on nuclear morphology. **A)** Representative snapshots showing the equilibrium morphologies of the nucleus, vacuole and cell at varying the vacuole enlargements (left to right), vacuole:nucleus bending rigidity, and +/- tethering. **B)** Relative nuclear size, **C)** nuclear displacement, and **D)** nuclear roundness as a function of vacuole size for cases with increasing vacuole bending rigidity and +tethering. **E)** Relative nuclear size, **F)** nuclear displacement, and **G)** nuclear roundness as a function of vacuole size for cases with decreasing vacuole bending rigidity and +tethering.

